# Genome-wide characterization of PRE-1 reveals a hidden evolutionary relationship between suidae and primates

**DOI:** 10.1101/025791

**Authors:** Hao Yu, Qingyan Wu, Jing Zhang, Ying Zhang, Chao Lu, Yunyun Cheng, Zhihui Zhao, Andreas Windemuth, Di Liu, Linlin Hao

## Abstract

We identified and characterized a free PRE-1 element inserted into the promoter region of the porcine *IGFBP7* gene whose integration mechanisms into the genome, including copy number, distribution preferences, capacity to exonize and phyloclustering pattern are similar to that of the primate Alu element. 98% of these PRE-1 elements also contain two conserved internal AluI restriction enzyme recognition sites, and the RNA structure of PRE1 can be folded into a two arms model like the Alu RNA structure. It is more surprising that the length of the PRE-1 fragments is nearly the same in 20 chromosomes and positively correlated to its fracture site frequency. All of these fracture sites are close to the mutation hot spots of PRE-1 families, and most of these hot spots are located in the non-complementary fragile regions of the PRE-1 RNA structure. Sequence homology analysis showed that the PRE-1 element seemed to share a common ancestor 7SL RNA with primates but was generated by different evolutionary model, which suggests that the suidae may be the closest relatives to primates in laurasiatheria.

## Introduction

Short interspersed nuclear elements (SINEs), a type of class 1 transposable element, have been promoted as valuable evolutionary markers in the population, species, genus and familial strata^1^. Since the mammalian radiation, the SINE family evolved differently in each lineage. All of the known SINEs are derived from tRNAs, with the exception of the primate Alu and rodent B1 families, which are derived from 7SL RNA (a component of the signal recognition particle)^2-8^. The Alu family was named for an internal AluI restriction enzyme recognition site. It descended from 7SL RNA 65 million years ago^9^ and propagated more than 1.2 million copies in the human genome, covering 11% of the genome’s total length, and is overrepresented in gene-rich/GC-rich regions. This propagation has resulted in the generation of a series of Alu subfamilies of different ages^10^; each primate genome has distinct subfamilies of the Alu element. Alu elements are specific to primates and only one type of SINE is found in the human genome. On the other hand, four distinct SINE families were found in the mouse genome: B1, B2, ID, and B4, which are unrelated to each other. B1 shares some sequence similarity with Alu and was thought to be a 7SL RNA-derived SINE. B1 and B2 elements occupy approximately 5% of the mouse genome, with approximately 550,000 and 350,000 copies, respectively^10^.

With a novel kind of 7SL RNA-derived SINEs, Tu types I and II have been identified as the Alu-like sequences in Scandentia (tree shrews)^11^ as well as other rodents in the recent years; all the 7SL RNA-derived SINEs have eventually been considered Supraprimate-specific^12,13^. The Alu elements (primate-specific SINEs) were initially considered to be selfish entities propagating in the host genome as “junk DNA”^14^. However, studies in the past decade have revealed that their insertion alters the structure and expression of genes, including influencing polyadenylation^15,16^, splicing^17-19^ and the adenosine deaminase that acts on RNA (ADAR) editing^20-23^. It now appears that there is a large reservoir of potential regulatory functions that have been actively participating in primate evolution, and the evolution of Alu subfamilies interacts in a complex way with other aspects of the whole genome. However, until now, the underlying evolutionary forces responsible for the origin and divergence of the SINEs remain obscure. The occurrence of 7SL RNA-derived SINEs was still thought to be a primate/Supraprimate-specific evolutionary event originating 65 million years ago because no new 7SL RNA-derived SINEs have been mined from other mammalian genomic data.

Although mammals share most of the same genes, the pig has always been thought to be a better biomedical model to humans than rodents due to its similar biochemical and physiological functions. The pig genome has been decoded, providing an necessary resource for mining similar genomic elements in non-coding regions. The SINE composition, being the second most abundant in terms of genome coverage^24^, was the first to be considered for genomic mining. In this report, we described a parallel world of porcine PRE-1 that almost copies all of the genomic performance of the human Alu element in the genome. In addition, we further explained that the formation of PRE-1 fragments of different lengths is based on a mutation hot spot-dependent fracture mechanism. Lastly, we compared the evolutionary behavior of porcine PRE-1 with that of human Alus and concluded that they are both descended from 7SL RNA and generated by different, independent evolutionary pathways. Therefore, all of the genomic evidence for a closer taxonomic relationship between the suidae and primate are not limited to the genes but also extended to the SINEs.

## Results

### Characterization of the PRE-1 element isolated from pig genome

The PRE1 element was first discovered as a polymorphic insertion in the 5’-flanking region, approximately 686∼985 bp upstream from the transcription initiator ATG codon of *IGFBP7* in our early experiments, as shown by agarose gel electrophoresis (Fig. 1a). Sequencing result of the PCR products revealed it is highly similar to the PRE-1 P27 reported earlier in Suidae^25^. (Fig. 1b), With a comprehensive structural and sequence conservation analysis, we found the element is a 299 bp long SINE inserted into the complementary strand (Fig. 1c) that is composed of two regions that are reverse complementary to each other (A-region: 1-115 bp, B-region: 116-210 bp) (Fig. 1d). The A region and B region share a 53-55 bp common homologous region with 63% similarity (Fig. 1e), each of which has an AluI restriction enzyme recognition site: AG/CT at the 3’end. The insert is terminated by a trinucleotide repeat (the main repeat unit is CAA) and a short poly(A) tail (Fig. 1e), which is a short intact direct repeat (GAATAACGGGCTTTT) derived from the site of insertion (blue arrows) (Fig. 1f).

**Figure 1.**
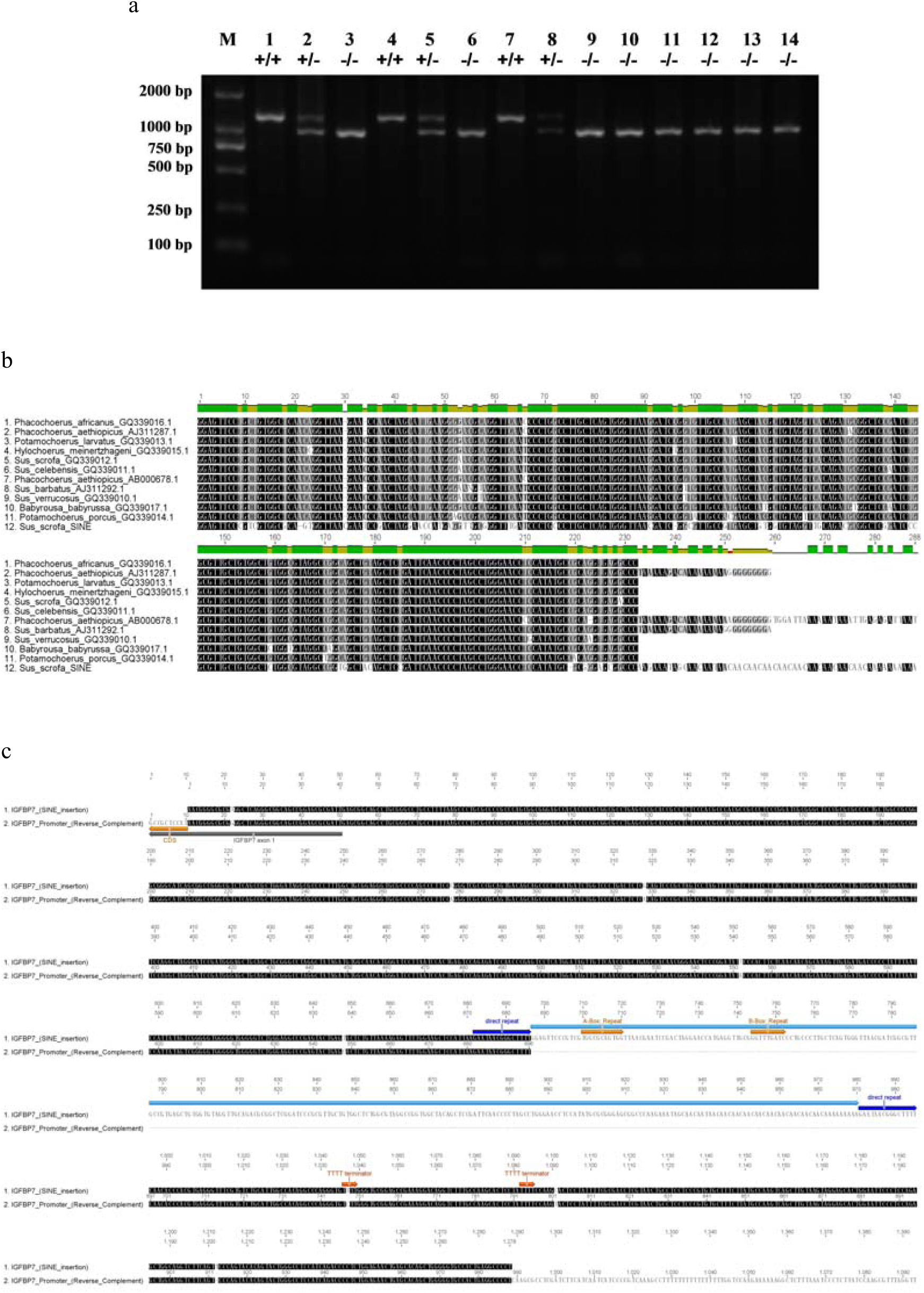
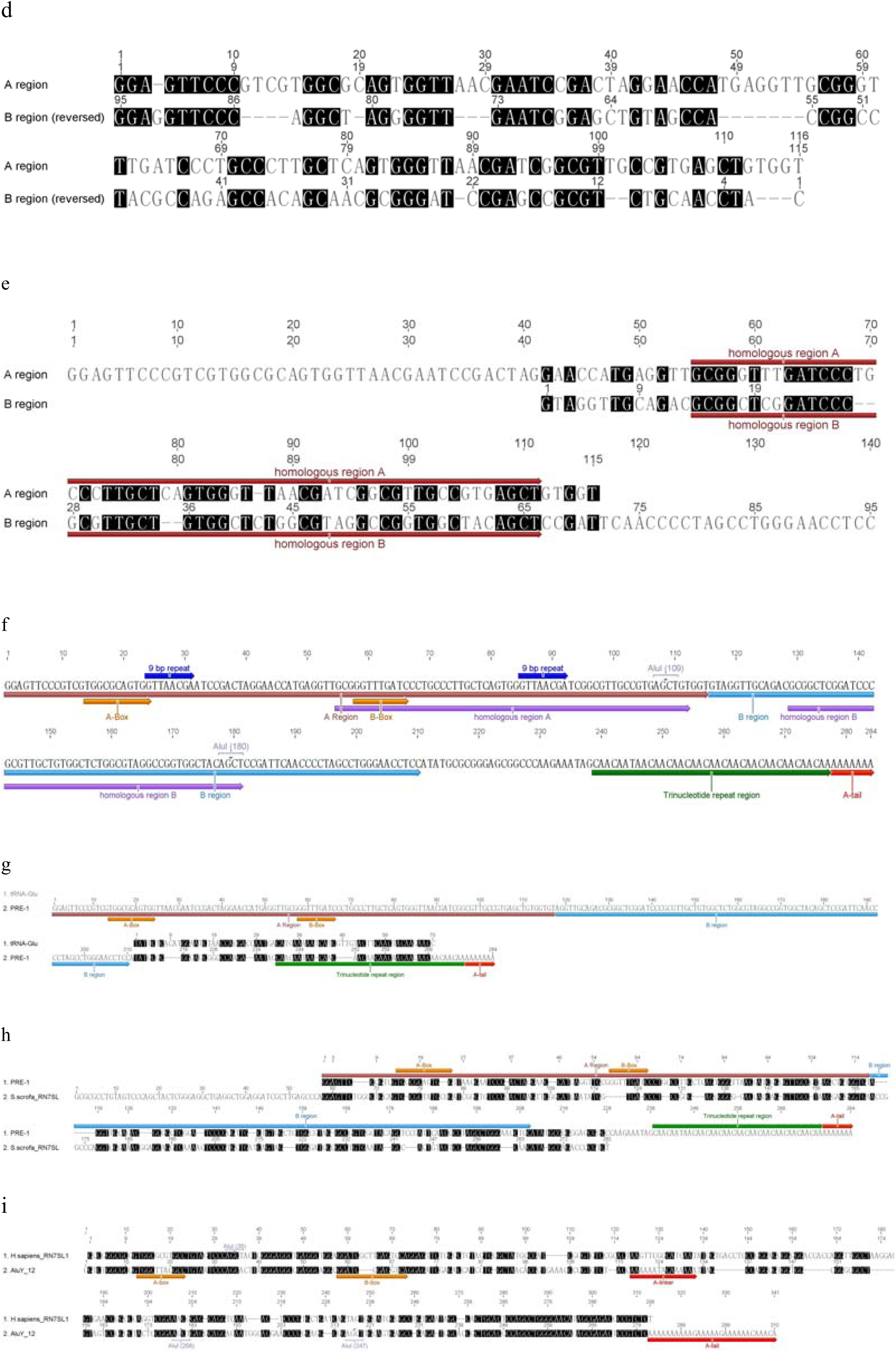
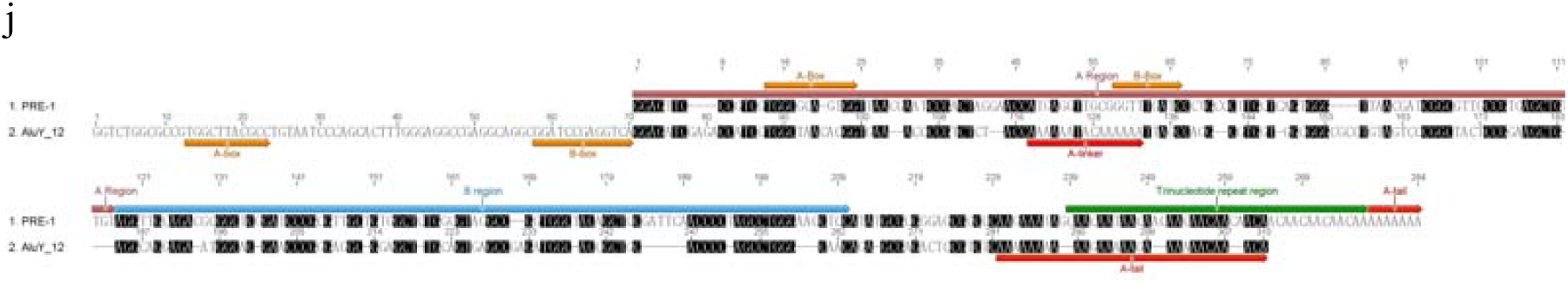
Characterization of the PRE-1 element isolated from pig genome. **(a)** The distribution of the insertion +/- polymorphism in the 5’-flanking region of *IGFBP7* in individuals from 7 pig breeds. (Lane M:2000 bp DNA marker, Lane 1-3: Tibetan pig, Lane 4-6: Wildboar in Northeast China, Lane 7-9: Min pig, Lane 10: Xiang pig, Lane 11: Yorkshire, Lane 12: landrace, Lane 13: Duroc, Lane1 4: Pietrain; -/- homozygous for the absence of the insertion; +/+ homozygous for the presence of the insertion; +/- heterozygous for the presence/absence of the insertion). **(b)** Alignment of SINE PRE-1 P27s in Suidaes with the PRE-1 element (IGFBP7) (c) Alignment of a sequenced DNA sample containing the insertion with the IGFBP7 promoter region sequence (NC_010450). This insertion sequence has two RNA-polymerase-III promoters (A and B boxes) but does not encode a terminator for RNA-polymerase-III. Instead, it seems to utilize the TTTT terminator downstream of the insertion site at various distances on IGFBP7 promoter to terminate transcription. **(d)** Alignment of the A region and the reverse-complement of the B region by Global Alignment (Needleman-Wunsch) for the PRE-1 element. **(e)** Alignment of the A and B regions by global alignment with free end gaps; 53-55 bp homologous regions with 63% similarity in the A and B regions were identified. **(f)** The architecture of the complete PRE-1 element with the direct repeat excluded. **(g)** Alignment of the PRE-1 element with porcine tRNA-Glu. (h) Alignment of the PRE-1 element with porcine 7SL RNA. **(i)** Alignment of the AluY_12 element with human 7SL RNA. The most notable feature of a representative primate Alu element is the presence of two “A-rich” regions.. The first “A-rich” region is near the center of the element with a consensus sequence of A5TACA6. It separates the sequence into two halves, left arm and right arm, and is often called the “A-linker”. The 5’ half of each sequence contains an RNA-polymerase-III promoter (A and B boxes). The second “A-rich” region is located 3’ of the right arm, 150 bp downstream of the first A-rich region, near the end of the Alu body, and almost always consists of a run of As that is only occasionally interspersed with other bases. It is often called the “A-tail”. **(j)** Alignment of the AluY_12 element with the PRE-1 element.

Postulating that this insertion sequence may be an Alu-like SINE, we further compared this insertion sequence with 40 representative primate Alu sequences selected from the AF-1 database (http://software.iiar.res.in/af1/index.html). AluY_12 shows the highest nucleotide similarity with the Alu-like sequence (Supplementary Fig. 1). We also aligned this Alu-like SINE with tRNA-Glu^26^ and the 7SL RNA gene of the pig (Fig. 1g, h), showing that just like the primate Alu element, this PRE-1 also has A- and B-boxes (internal RNA polymerase III promoter) but with two 9 bp repeats (GGTTAACGA) in the A region. From the alignment of AluY_12 with human 7SL RNA (Fig. 1g), we see that in humans, the original A- and B-boxes from 7SL RNA were lost and a new set of A- and B-boxes evolved; however, in this PRE-1 element, the A- and B-boxes perfectly match the canonical sequence found in tRNA genes. The sequence of primate Alu A- and B-boxes slightly diverges from the canonical A-and B-box sequences, TRGYnnAnnnG (11 nt) and GWTCRAnnC (9 nt), which are the hemiascomycetous signatures of the Pol III transcriptional promoters for tDNA. No A-rich linker was identified between the A region and B region in this PRE-1 element, but it has more sequence similarity to the 7SL RNA central region than is found in the human Alu element.

### The copy number of the PRE-1 repeats in pig genome

To identify more PRE-1 elements, we used BLASTn to search “somewhat similar sequences” to this first identified PRE-1 element against the database of the NCBI Pig Genome, and 1037475 blast hits (length:12-350bp) of the PRE-1 repeat were found on both strands in the genome, which together account for nearly 8.21% of the sequenced region of the pig genome. The correlation coefficient between the copy number of PRE-1s and the chromosome length is 0.973 (Fig. 2a), large chromosomes were necessarily associated with more PRE-1 repeats.

**Figure 2.**
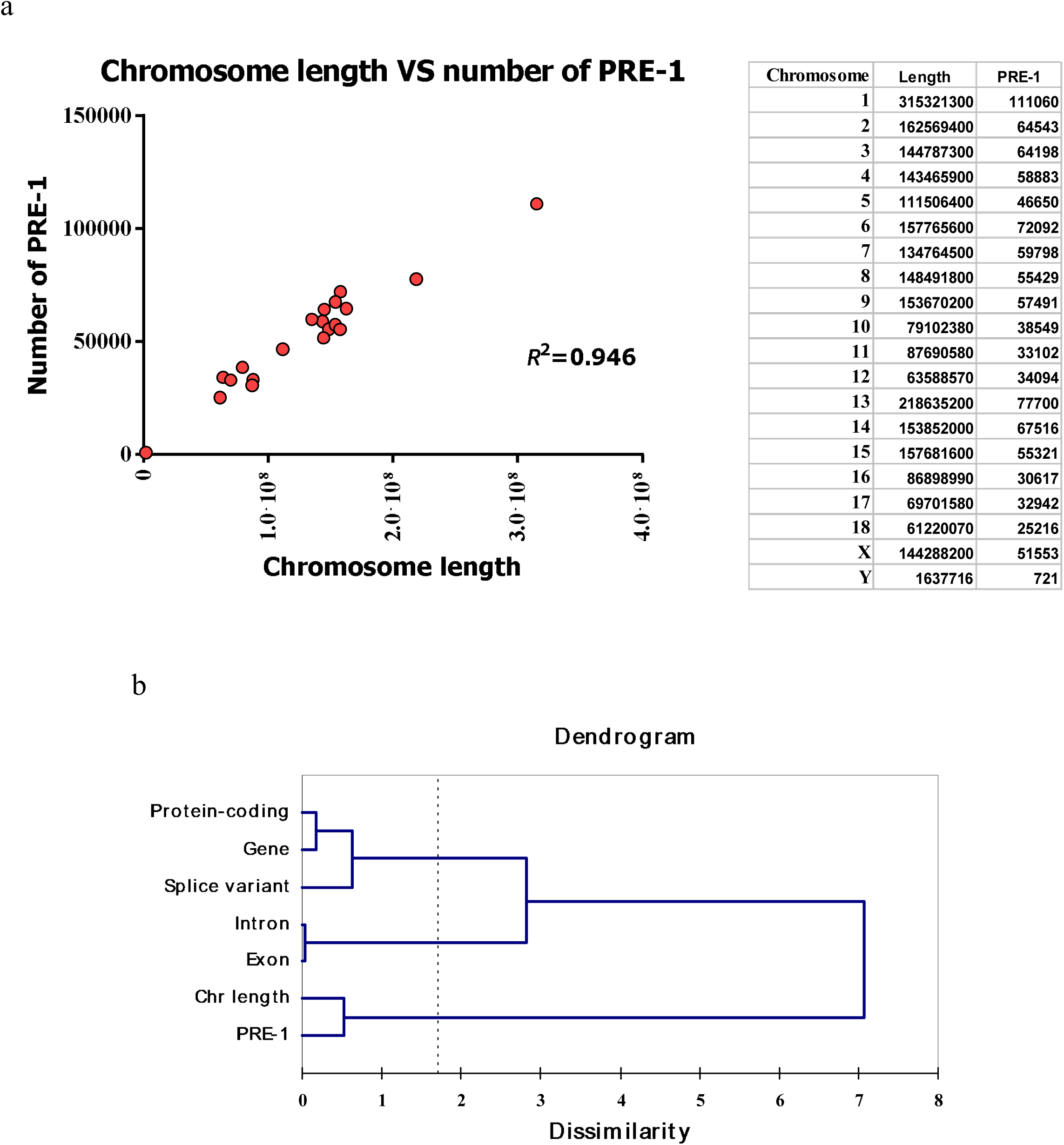
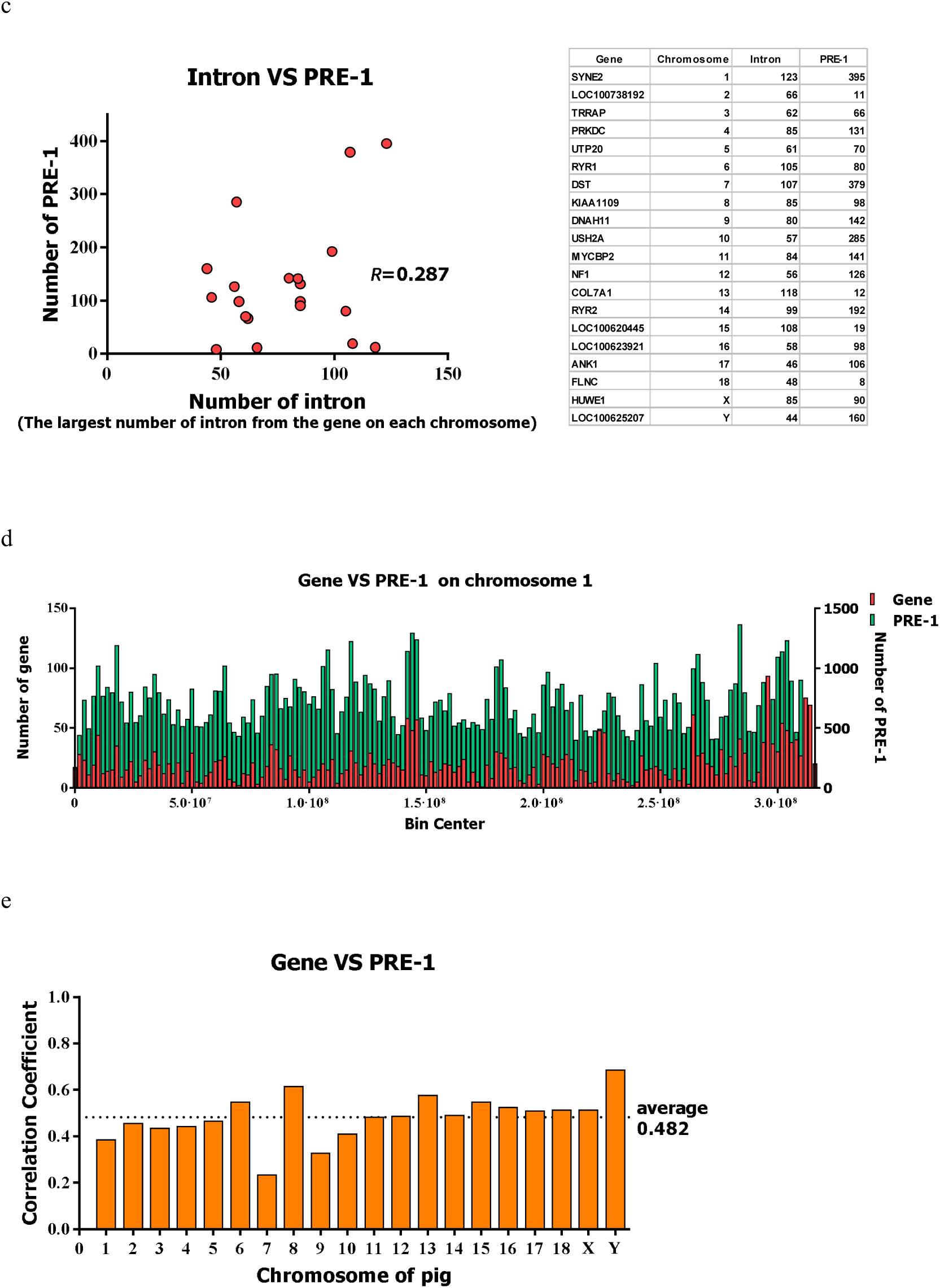
The copy number of the PRE-1 repeats in the pig genome. **(a)** Variation of PRE-1 repeats and chromosome lengths in pig me. **(b)** Ward’s dendrogram of Hierarchical Cluster Analysis using 7 variables **(c)** The correlation of the number of PRE-1s and intr ithin the gene with the largest number of exons in each pig chromosome.. **(d)** The comparison of density distributions between the P -1 repeats and genes on pig chromosome 1. **(e)** The correlation coefficients of density distributions between the PRE-1 repeats and s on 20 pig chromosomes.

Additionally, we performed a cluster analysis to investigate the correlation among the number of PRE-1 repeats with other important genomic features (numbers of genes, protein-coding sequences, splice variants, exons, introns, and the length of chromosome); all these data are grouped in terms of the chromosome, and Ward hierarchical clustering was applied (Supplementary Table. 1).

The dendrogram revealed that the number of PRE-1 repeats is correlated with six variables in the following order of correlation strength: chromosome length > exons > introns > splice variants > genes > protein-coding sequences. (Fig. 2b), so we selected a small sample including the gene that is composed of the most exons in each chromosome and calculated that the correlation coefficient of the numbers of introns and PRE-1s in each gene region is only 0.287 (Fig. 2c). As primate Alu elements accumulate preferentially in gene-rich regions, we further compared the density distribution of genes and PRE-1 elements for 20 chromosomes, where each chromosome was split into 1000 kb intervals and PRE-1 size and gene size was calculated individually for all of these regions. We found that the overall trend of the density distribution between of the two is very similar, with some exceptions in a few areas in chromosome 1;, the average correlation coefficient of 20 chromosomes is 0.482. (Fig. 2d, 2e, Supplementary Table 2, 3).

We deduced that gene enrichment in a genomic region is likely to attract more PRE-1s parasited in the introns, but the genomic feature in the gene-rich region, such as the intron size instead of intron number, may be a necessary condition for accommodating more repeats.

### The ength-frequency distributions of PRE-1 repeats

A full-length PRE-1 is approximately 300 nucleotides in length, but the results from the PRE-1 BLASTn search showed that the dynamic size range of these “somewhat similar sequences” is approximately 12 -350 bp. We calculated the length-frequencies of PRE-1 repeats on both strands of 20 chromosomes. Surprisingly, not only the total numbers, but the length-frequency distributions of sense and antisense PRE-1 elements are highly similar on each chromosome. When plotting the positive Y-value of the length-frequency of sense PRE-1 elements and negative Y-value of antisense PRE-1 elements from 10-350 bp at 10 bp segments along the X-axis, the length-frequency distributions of sense and antisense PRE-1 elements mirrored each other vertically, thus presenting a perfect symmetrical “tadpole” type curve (Fig. 3a). Only the Y chromosome is limited by insufficient PRE-1 repeats(Fig. 3b), all the other 19 chromosomes follow the same rule (Supplementary Fig. 2, Table. 4).

**Figure 3.**
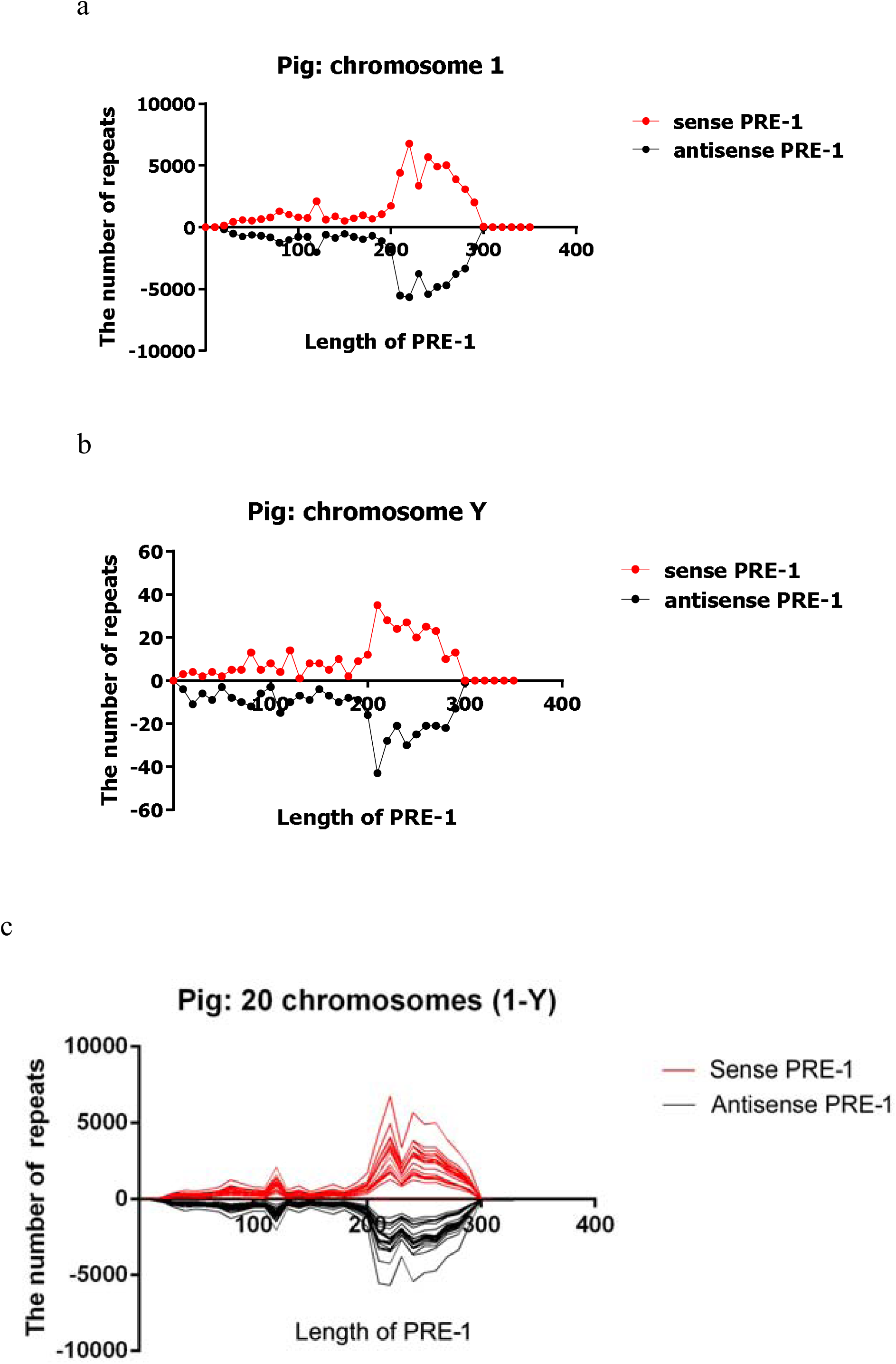
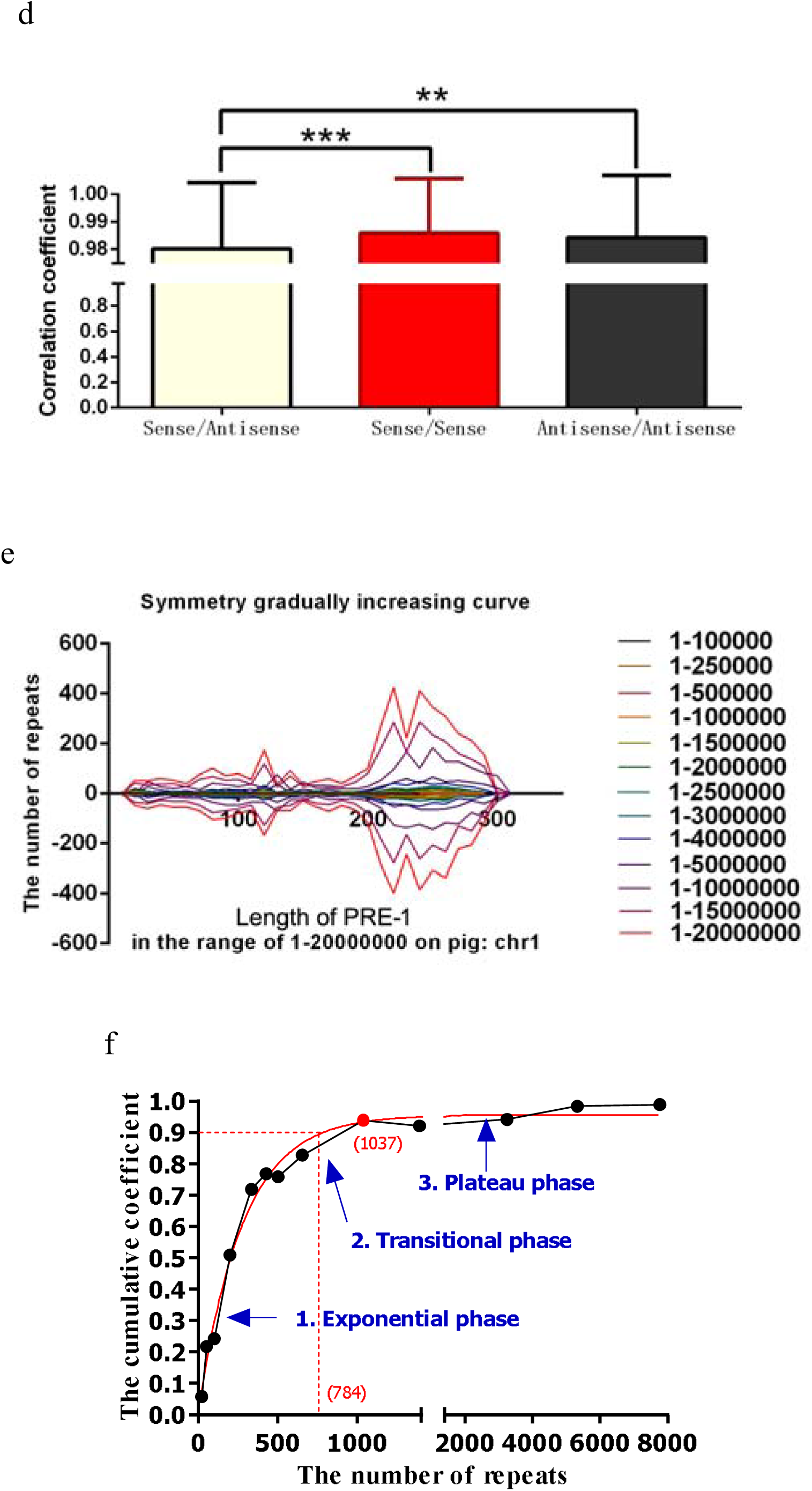
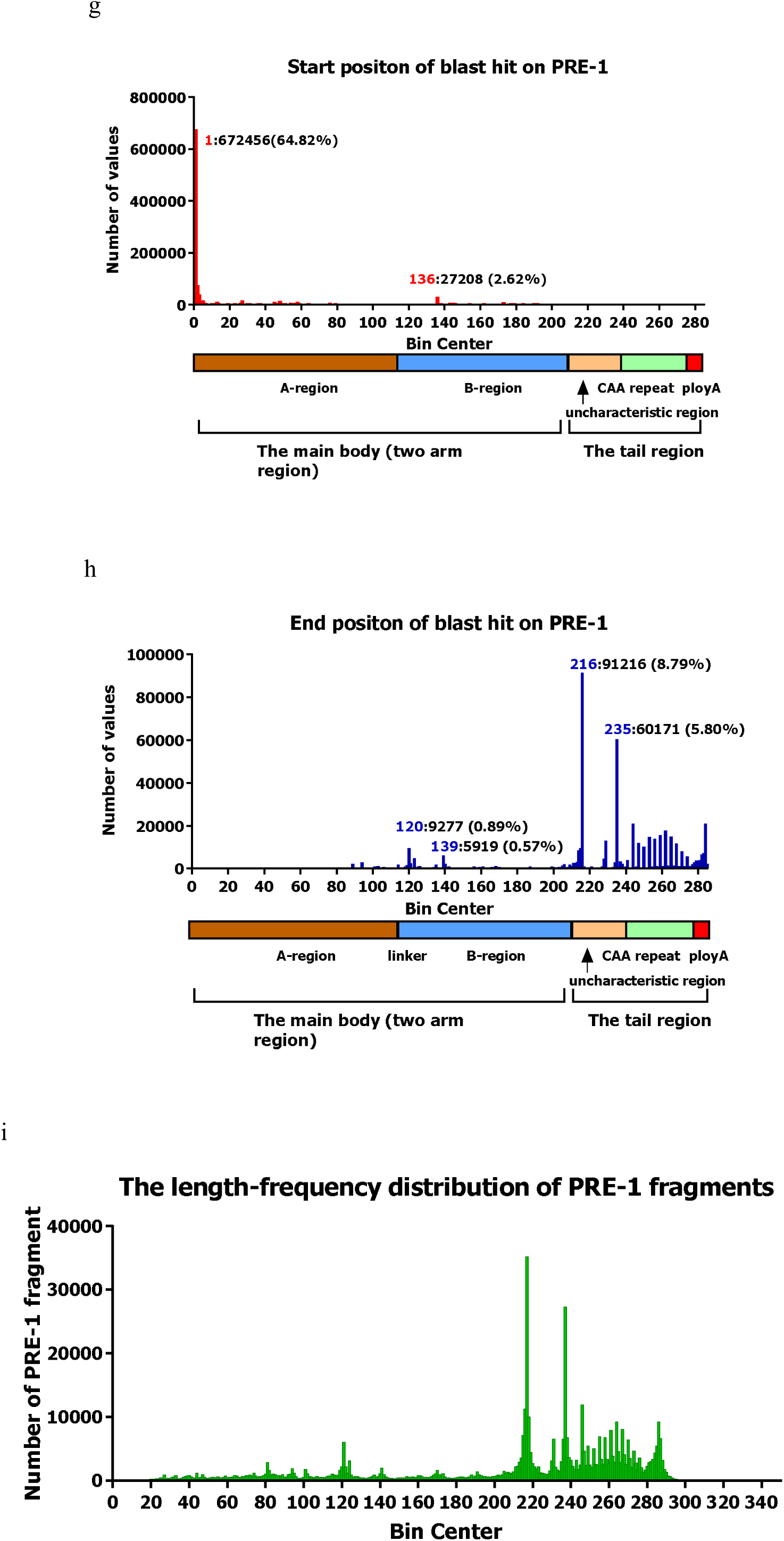
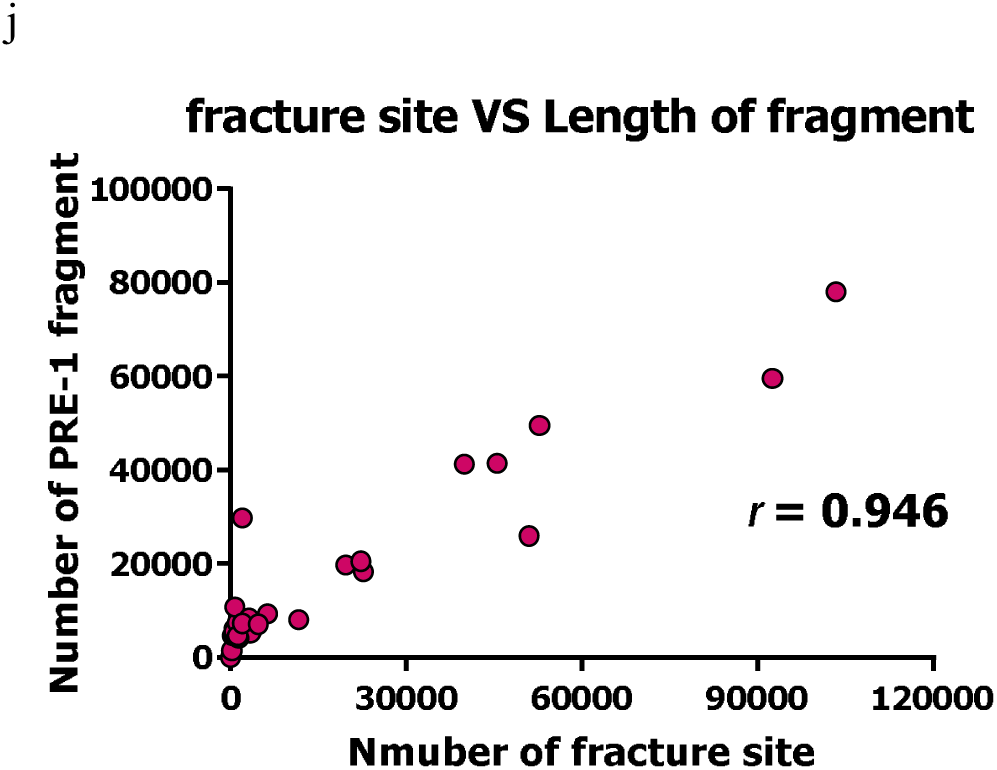
The length-frequ stributions of PRE-1 repeats in the porcine genome. (a, b, c) The symmetrical distribution of various lengths in sense and ense PRE-1 repeats on pig chromosome (a) 1, (b) Y, and (c) all 20 chromosomes. (d) Symmetry differences between sense/sen RE-1, antisense/antisense PRE-1, and sense/antisense PRE-1 on 20 chromosomes. Paired t-tests indicated that the correlation co ient of sense/sense PRE-1 and antisense/antisense PRE-1 was higher than that of sense/antisense PRE-1 (for the one-tailed test, 0.01,*** p<0.001). (e) Curves of the sense/antisense PRE-1 distribution. With the length size increased in the range: 1-20000 bp on pig chromosome 1, more of the repeats are amplified, increasing the symmetry of the length-frequency distribution of E-1 repeats. (f) Growth curve of the Pearson coefficient in the region ranging from 1 to 20000000 bp on chromosome 1. The number PRE-1 repeats increased from 24 to 7768. The values of cumulative Pearson coefficients of the length-frequency distribution be sense and antisense PRE-1 elements grow faster as the PRE-1 sample size increased in the range of 1-1000. Limiting factor ow and eventually stop coefficient growth. The coefficient seemed to follow a growth curve with three phases: exponential phase, tran ional phase, and plateau phase (Y= -1186+(1186.956)/(1+10^(-0.001571X)), R2= 0.9884). (g) The integration initiation site frequen istribution of PRE-1 fragments on the whole genome (h) The integration termination site frequency distribution of PRE-1 fragments the whole genome. (i) The length-frequency distribution of PRE-1 fragments on the whole genome (j) The correlation of frequency nu r between fracture sites and fragment lengths on the whole pig genome (The interval of frequency distribution for fracture sites and fragment lengths is 10 bp).

When integrating the PRE-1 length-frequency distribution symmetrical curves of the 20 chromosomes into one axis, two strong correlations of length frequency distribution curve were clearly observed. The correlation of sense/sense PRE-1s (red line) or antisense/antisense PRE-1s (black line) strands among the 20 chromosomes is higher than that of sense/antisense PRE-1s on each chromosome (Fig3c). Paired t-tests were performed to compare the difference between the correlation coefficients of sense/sense PRE-1s and antisense/antisense PRE-1s versus sense/antisense PRE-1s (control group) (Fig. 3d, Supplementary Table.5).

For investigating how the copy number of PRE-1s affect the degree of symmetry of the length-frequency distribution curve, we selected 13 pairs of data (calculating the correlation coefficients corresponding to the number of PRE-1 elements) by an increasing interval step in a region ranging from 1 to 20000000 bp on chromosome 1 and plotted a growth curve of the correlation coefficient (Fig. 3e). By performing non-linear regression analysis, the 4 parameter logistic non-linear regression model had been tested to fit the curve well. According to this non-linear regression model, we deduced that when the total number of PRE-1 elements cumulated to approximately 784, the Pearson coefficient will reach 0.9; when the number of PRE-1 elements increased to 1000, the Pearson coefficient will plateau at 0.92-1 (Fig. 3f, Supplementary Table 6).

According to the regression equation, we know that the number of PRE-1 elements is only 721 on the Y chromosome, so the correlation coefficient is only 0.902, which is short of the plateau and explains the lack of high symmetry of the curve.

We further calculated the percentage of integrating and break sites of these repeats initiated from the complete PRE-1 elements on all 20 chromosomes by the hit location information extracted from the results of the BLASTn search. In total 1037475 PRE-1 blast hits were used, and the proportion of integrating sites of the PRE-1 blast hits was displayed in a plot chart as shown in Fig. 3g We found that 64.82% of the integrating sites in the PRE-1s are initiated from position 1 and that 91.45% of them are from the A region; there is only one significant integration site in the B region at position 136 (Fig. 3 g). Meanwhile, 42% of the fracture sites are concentrated in the tail region. Positions 216 and 235 are located in the uncharacteristic region and have the highest fracture probabilities, accounting for 8.79% and 5.80%, respectively, but in the trinucleotide repeat region, the fracture probability showed a normal distribution (Fig. 3 h). There are two fracture hotspots in the B region, one is at position 120 (0.89%) and the other is at position 139 (0.57%). The frequency distribution of fracture sites and fragment lengths showed a highly positive correlation, with a 0.946 correlation coefficient (Fig. 3i, 3j).

### Exonization of the PRE-1 elements in pig genome

Although Alu elements are ubiquitously interspersed in gene-rich regions throughout the human genome, with the majority located in introns, most Alu elements have remained silent, whereas a small subset have undergone exonization. More than 5% of the alternatively spliced internal exons in the human genome are derived from Alu elements^27^. Compared to normal cells, Alu exonization occurred in one of the non-coding transcripts or in most leukemia/cancerous cells at a very high level^28^. The exonization of Alu sequences has been shown to be regulated not only by splice sites but also by splicing regulatory sequences within the Alu elements^29^.

In total, 2337 PRE-1 blast hits were found on the 1913 transcripts, including 673 splice variants from 1240 genes found by BLASTn search against the NCBI reference RNA sequence of Sus scrofa, consisting of 311 non-coding RNAs (accounting for 18% of total non-coding RNAs) and 929 mRNAs (accounting for 4% of total coding RNAs) (Fig. 4a).

**Figure 4.**
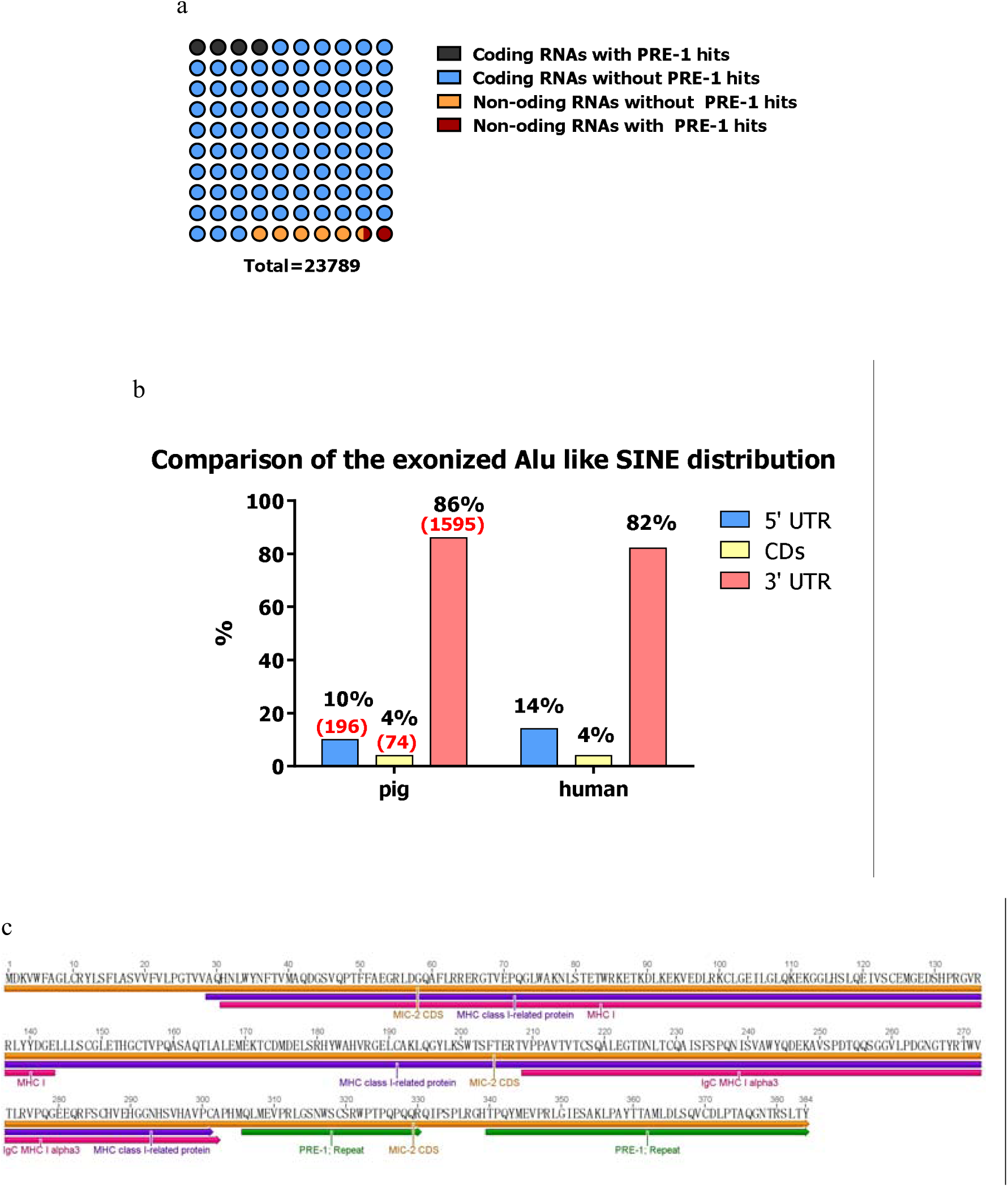
Exonization of the PRE-1 repeats in pig genome. (a) Proportion of PRE-1 hits in the total transcribed RNAs. (b) Comparison of the Alu-like SINE distribution ratio on mRNA between pig and human (c) Two fragments of PRE-1 repeats are integrated into the coding region of MIC-2

We found 196 PRE-1 blast hits in the 5’UTR of mRNAs, 1595 PRE-1 blast hits in the 3’UTR of mRNAs, and 74 PRE-1 blast hits in the protein-coding regions. These results indicated that the 3’UTR of mRNAs that contain PRE-1 repeats are predominantly similar to the distribution on the human Alu element (Fig. 4b)^30^, although most of the PRE-1 blast hits in the protein-coding regions are less than 100 bp and adjacent to the coding region boundary. A few large fragments of PRE-1 elements were found integrated in coding regions and involved in the formation of new proteins such as MHC class I related antigen 2 (MIC-2) (Fig. 4c).

### Clustering and subfamily relationships of the PRE-1 family

Human Alu repeats can be divided into at least three^31^ and probably four^32^ subfamilies that are apparently of different ages. To clarify the evolutionary relationships of PRE-1s, we performed a local comparison of the PRE-1 sequences derived from medium-gene density regions of each chromosome for the proportion of “embed PRE-1s” and “free PRE-1s”. Accounting for the complete structural features of the PRE-1 elements, we assume that the majority of 280-350 bp insertions arise as a result of new retrotransposition events. We separated a small dataset containing 517 sequences screened from 8787 PRE-1 sequences, and further optimized by MaxAlign (http://www.cbs.dtu.dk/services/MaxAlign/). In total, 347 sequences were used to construct a Maximum-likelihood tree, including the PRE-1 elements we have identified before, PRE-1 P27 and 7SL RNA (rooted as an outgroup). The phylogenetic analysis of PRE-1 repeats derived from the unbiased small sample reveals three different categories of repeats (PRE-1_A, PRE-1_B and PRE-1_C), meaning at least three propagation waves occurred in the evolutionary process of PRE-1 elements. PRE-1_B and PRE-1_C families both contain 10 subclades. The sus scrofa PRE-1 P27 belong to the PRE-1_B family, whereas the PRE-1 we identified was a young element from the PRE-1_C family, of which a more abundant set of young elements are included; there have been 56.2% more PRE-1 retrotransposition events as a result of the expansion of the PRE-1_C family (Fig. 5a)

**Figure 5.**
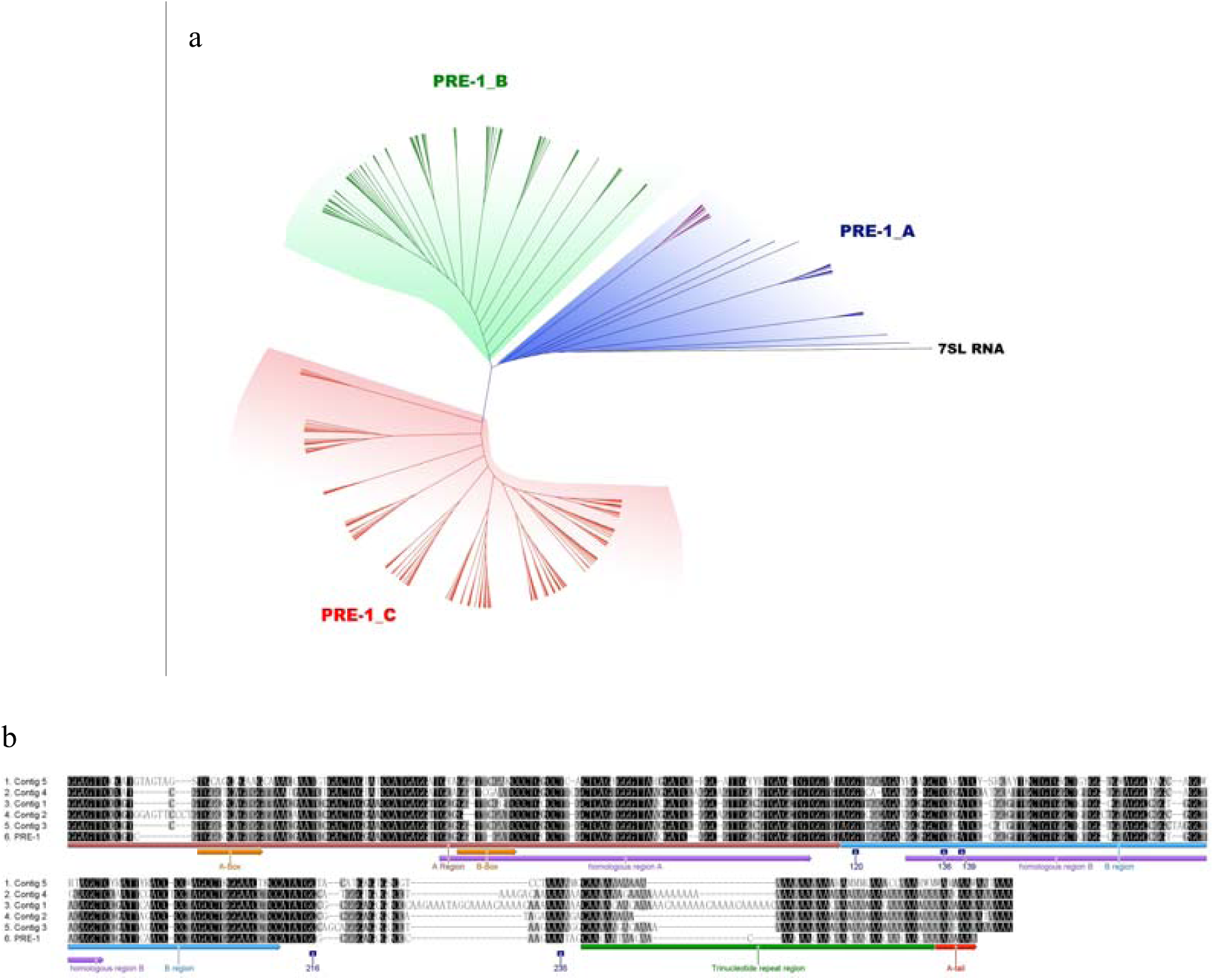
(a) Maximum-likelihood tree based on 349 PRE-1s (PRE-1 (IGFBP7), PRE-1 P27 included) and 7SL RNA. **(b)** Comparison of a multiple alignment of 5 consensus sequences with a standard complete **PRE-1**, the integration initiation site and termination site are marked deep blue.

We assembled all of these 347 PRE-1 sequences into 5 consensus sequences using DNA Dragon. By aligning these sequences, we found that each integration initiation site or termination site is located ahead of a hypervariable region in the multiple alignment; three mutation regions with the largest degree of variation are all linked with sites: position 216, position 235 and the trinucleotide repeat region. (Fig. 5b).

We found 98% of the 347 PRE-1 elements contain two conserved AluI restriction sites by performing restriction enzyme analysis.

## Discussion

Unlike all other transposons, most SINE families do not share a common origin and were independently generated in different host lineages from cellular and LINE modules^12^. The specific SINEs or SINE families are usually restricted to respective orders, rarely crossing order boundaries^33^. In recent years, a novel kind of 7SL RNA-derived SINE, Tu type I, had been identified to be a 7SL RNA-derived SINE in scandentians (tree shrews)^11^. These 7SL RNA-derived SINEs have already been considered a supraprimates-specific SINEs^13^. The 5’end of SINEs are derived from 7SL RNA (Alu, B1, and their derivatives in primates, rodents, and tree shrews), the origin of the central part of most SINE families is obscure, and the 3’ end of SINEs descend from the 3’ end of the partner LINE^34^.

The first PRE-1 element was identified in the genome of the miniature swine by Singer et al. in 1987^35^ and was thought to be an arginine-tRNA gene^36^, which diversified at least 43.2 million years ago^37^. Approximately 98% of PRE-1 elements contain two conserved AluI restriction sites, (Supplementary Fig. 4), but only 40% Alu representative sequences in the AF-1 database have conserved double-AluI recognition sites in the B arm region, and the evolution of the PRE-1 element from 7SL RNA is more evident than that for the Alu element (Fig. 6a). The conclusion from our results is that the PRE-1 element maybe a unique SINE in the pig genome. It is more like a 7SL RNA-derived SINE than a tRNA-derived SINE. In the process of integration into the genome PRE-1 will fracture into fragments. As 7SL RNA was generated from tRNA, these small indirect tRNA derived fragments are usually considered to be mistakenly derived from tRNA.

**Figure 6.**
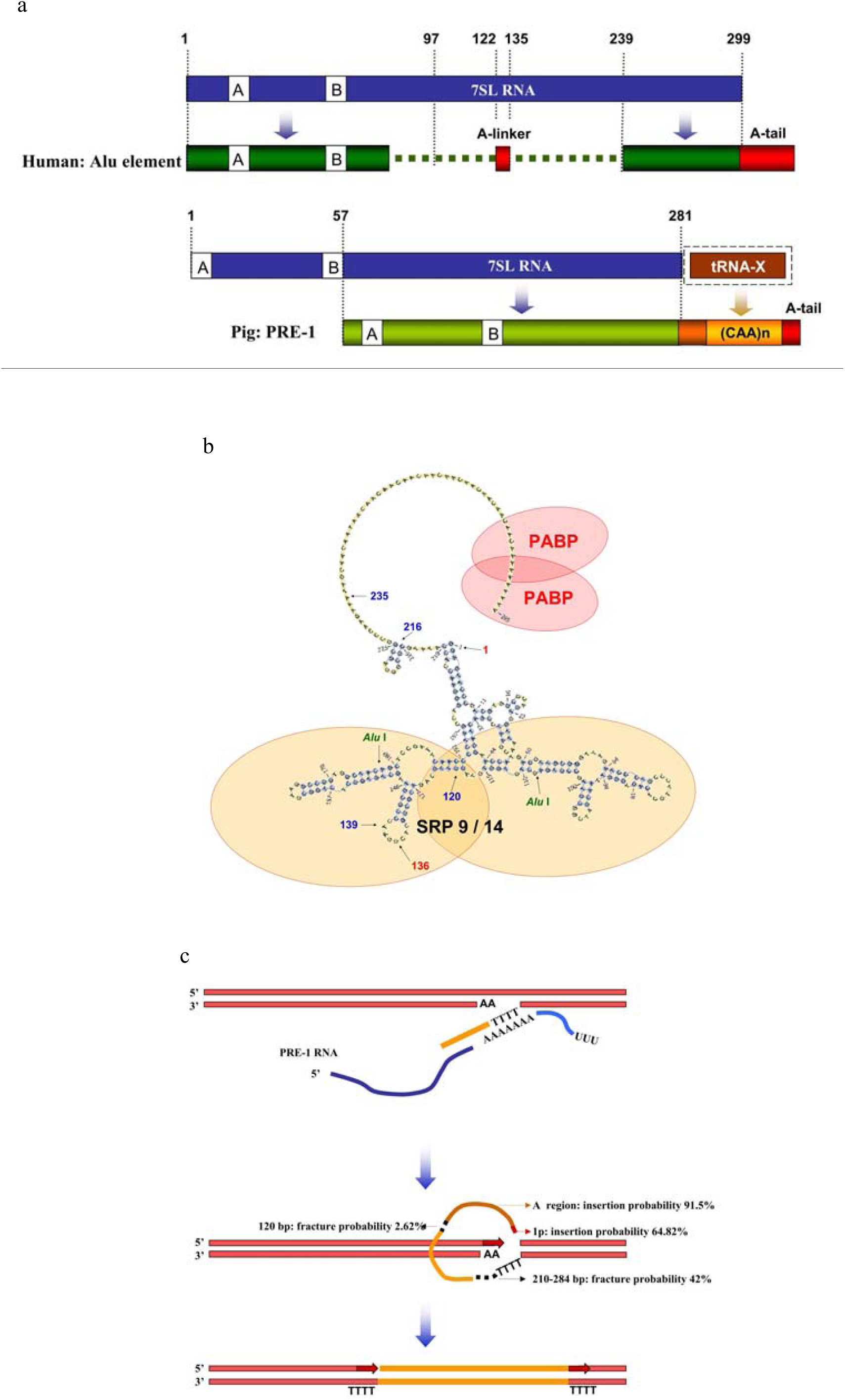
Birth and amplification mechanism of PRE-1 element. **(a)** The 7SL RNA-derived SINEs of humans and pigs probably arose by two different evolutionary models. Primate-specific SINEs evolved from a 7SL RNA sequence and gave rise to the part of Alu SINEs by a fusion of the 5’ and 3’ ends of the 7SL RNA gene with an obscure middle part. Independently, suidaes-specific PRE-1 elements seemed to evolve from a 7SL RNA gene by losing 57-bp on the 5’ end and hybridizing with tRNA (Glu or others) to complete the tail region. (b) Secondary structure of PRE-1 RNA. The PRE-1 RNA is predicted to fold into separate arms for each region (Supplementary Fig. 5). The two arms are suspected to bind the 7SL RNA SRP9 and 14 heterodimer, with an A-tail for polyA-binding protein (PABP) binding. (c) The target-primed reverse transcription mechanism (TPRT) of PRE-1. PRE-1 retrotransposition is also likely to occur through TPRT using the L1 retrotransposition machinery. The first nick at the site of insertion is often made by the L1 ORF2 protein’s endonuclease at the TTAAAA consensus site. The T-rich region primes reverse transcription by L1 ORF2 proteins on the 3’ A-tail region of the PRE-1 element. This creates a cDNA copy of the complete PRE-1 RNA. The mechanism for making the second nick on the other strand and integrating the other end of the PRE-1 coincides with losing part of the sequence at the fracture hot spot. A new set of short direct repeats (red arrows) is created during the insertion of the new PRE-1 element. During the TPRT process, PRE-1 elements will fracture and formed repeats of various lengths integrated in the pig genome.

Just like human Alu elements, PRE-1 elements tend to be clustered in the introns of gene-rich regions, so we abandoned the optimal interval of each chromosome to calculate the correlation instead of a unified interval length (1000 kb), concluding that the average correlation coefficient of the density distributions of genes and PRE-1 elements for all 20 chromosomes is only 0.482. As we observed on the COL7A1 gene, which has 119 exons on chromosome 13, most introns are extruded into “narrow introns,” within which 12 PRE-1 elements are accommodated, so we concluded that the numbers of “wide introns” may play a key role to explaining why PRE-1 elements are parasitic in gene-rich regions.

Although the PRE-1s propagated the nearly same number of repeat on both sense and antisense strands, and sensitive between them, but the length distribution of PRE-1 fragments is most intriguing and indicates that PRE-1 propagation is a continuous fragmented insertion process. The amplification mechanism of PRE-1 repeats is very similar to that of the primate Alu repeats, the variation of repeat size is because any position of the complete replicated strand can be inserted into the target site using a nick at its genomic integration site to allow target-primed reverse transcription (TPRT) to occur. During the insertion process, 64.8% of all PRE-1 elements can integrate into the genome from the initial site at position 1, 78% of all insert hotspots are concentrated in the range of positions 1-10, and 91.5% of the integrating sites are distributed in the A region. In the B region, there is only one insertion hotspot at position 136, which occupied 2.62% of all insert sites. Fractures occur along with insertions, and only 2.9% of all PRE-1 elements can retain full length (≥285 bp); 42% of all fracture sites are distributed in the tail region.

Positions 216 and 235 are the first (8.79%) and second (5.80%) most frequent fracture hotspots, respectively. The most abundant fracture hotspots occur in the trinucleotide repeat region ranging from positions 240-275, accounting for approximately 15.93% of all fracture sites. The trinucleotide repeat region formed a normal distribution of fracture sites, which matched the results of the main peak of the distribution of SINE length located approximately 263 bp as revealed by Xiaodong Fang et al^26^. from the WZSP genome. There are two significant fracture hotspots in the B region, one is in position 120 near the linkage of the A region and B region and comprises 0.89% of all fracture sites; the other is in position 139, comprising 0.57% of all fracture sites. According to the alignment of 347 complete PRE-1 elements, we found that all fracture hotspots are followed by a mutation region; it seemed that a higher degree of variation indicated a higher probability of fracture.

We further predicted the secondary structure of PRE-1 RNA using RNAstructure Webservers (http://rna.urmc.rochester.edu/RNAstructureWeb/Servers/Predict1/Predict1.html), The modle was chose for it’s the most minimal energy value shows that even without an A-linker, the structure of PRE-1 RNA can still be folded into a two-arm model where most mutations in the tail region are located in the non-complementary single-stranded region, which is the most fragile region of the PRE-1 RNA structure. The PRE-1 RNA structure further revealed that it evolved from 7SL RNA as it shares similarity to the primate Alu element based on the RNA structure. The PRE-1 RNA maybe also transcribed from the adjacent genomic site and is thought to assemble into a SRP9/14 heterodimer, with polyA-binding protein (PABP) and at least one other unidentified protein binding to the RNA structure. The SRP9/14 proteins and PABP help the PRE-1 RNA associate with a ribosome to associate with ORF2. PRE-1 RNAs then utilize the purloined ORF2 protein to copy themselves at a new genomic site using a process termed target-primed reverse transcription^23^. and now, we speculated that the gene cluster seems has dual effects of attraction and insulation on the PRE-1 or other SINEs during the integration process, the number of PRE-1 inhaled in the non-coding region of the gene region is significantly higher than that of in the non-gene regions, We are not sure about how long the fragment is at some insertion site, but only get the probability of the fragment.

In the past, we were always much more concerned with the evolution of genes rather than with other genomic elements and used genes as the most important markers to trace phylogeny, regardless of the fact that the genome was coevolved by the gene together with other genomic elements. In our research, the genomic performance of PRE-1 in terms of 7SL RNA-derived SINEs seemed convincing enough to classify the suidae into a family mainly inhabited by primates, so the divergence time of 7SL RNA evolutionary events may be re-adjusted back to 80-100 million years ago^38^, before the boreoeutherians diversified into Laurasiatheria and Euarchontoglires^39^. 7SL RNA began to diversify or hybridize from different tRNAs to form a variety of 7SL RNA-like SINE in boreoeutherian ancestors. One branch was found in the ancestor of primates and evolved in supraprimates; the other was found in the ancestor of suidaes and evolved in laurasiatherias. The similarity of PRE-1 and Alu elements further revealed a hidden kinship, such that suidaes are more closely related to the suraprimate/primate than any other laurasiatherias based on the 7SL RNA derived SINE composition of their genomes.

## Acknowledgments

We thank the Swine Genome Consortium for providing complete sequences and detailed annotations on the reference genome build 10.2 and the anonymous reviewers who helped improve this article. This research was supported by National Modern Agricultural Industrial Technological System (CARS-36), China Postdoctoral Science Foundation, National Natural Science Foundation Of China (Grant no. 31101781)

## Methods

The numbers and positions of PRE-1 repeats in complete genome were identified using BLASTn (http://blast.ncbi.nlm.nih.gov/Blast.cgi). Information about the associated genes, protein coding sequences, introns, exons, and splice variants of pig genome was retrieved from the annotations of each chromosome available on NCBI’s gene database.

Correlation between PRE-1 repeats and gene density was calculated for non-overlapping windows along the whole chromosome at 1000 kb intervals in term of chromosome size. The total PRE-1 size (base pairs in an interval occupied by a PRE-1 elements) was taken as measures of PRE-1 density, rather than its numbers.

Statistical analyses were performed using Excel, XLSTAT, and Graphpad Prism 6 software packages. Sequence alignments and graphic displays were generated using the Geneious Pro software package (version 4.8.3). The phylogenetic tree was constructed using the Maximum Likelihood method in MEGA (V.6.06), and the PRE-1 contigs were assembled by DNA Dragon 1.6.0 with 80% identity of complete overlapping fragments.

## Supplementary information

**Fig. 1.**
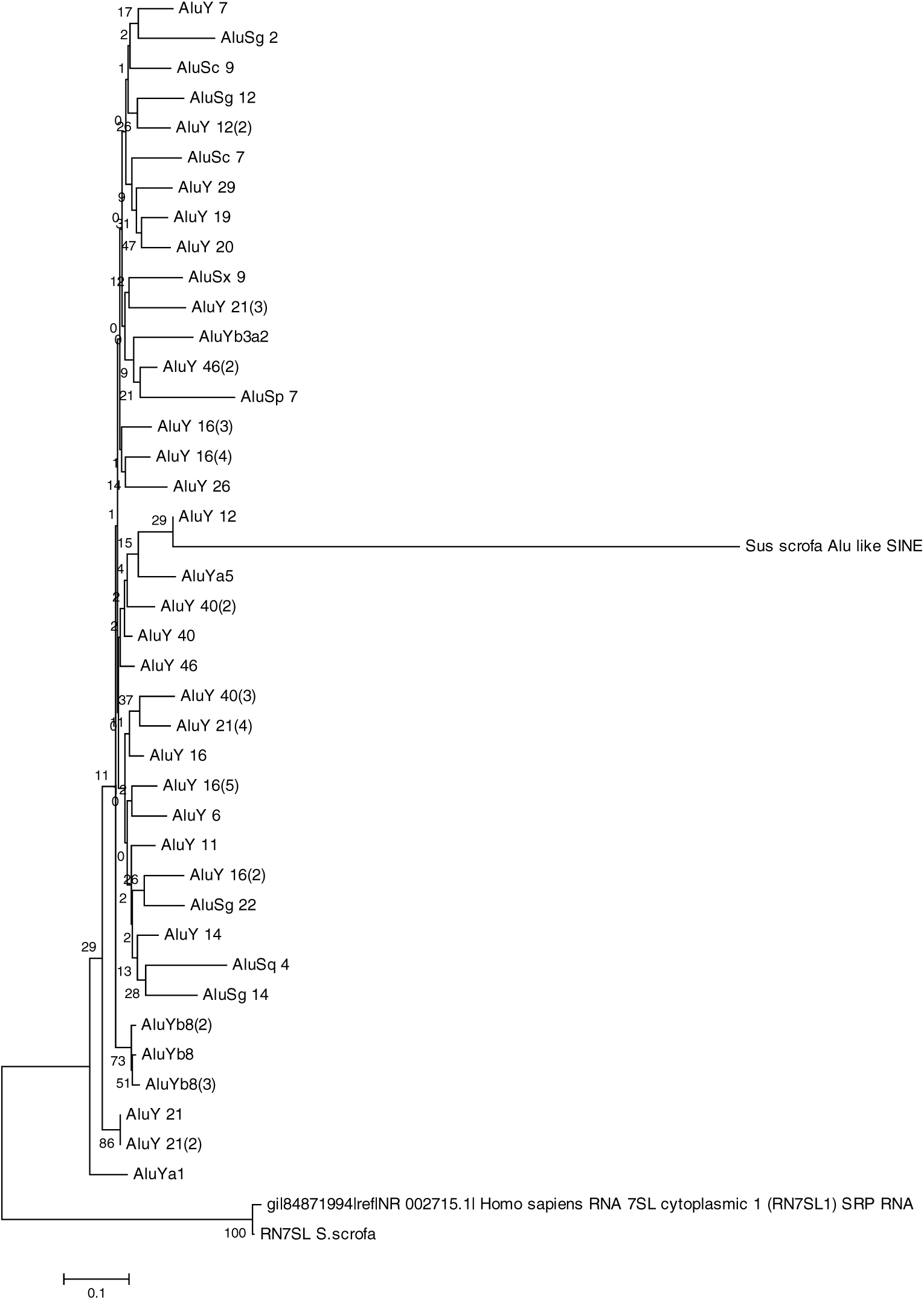
Clustering tree of porcine Alu-like SINE/PRE-1 with representative primate Alu sequences. A maximum likelihood tree produced by MEGA 6.06 using 40 primate Alu sequences from the AF-1 database with porcine Alu-like SINE/PRE-1 and 7SL RNAs of pig and human.

**Fig. 2.**
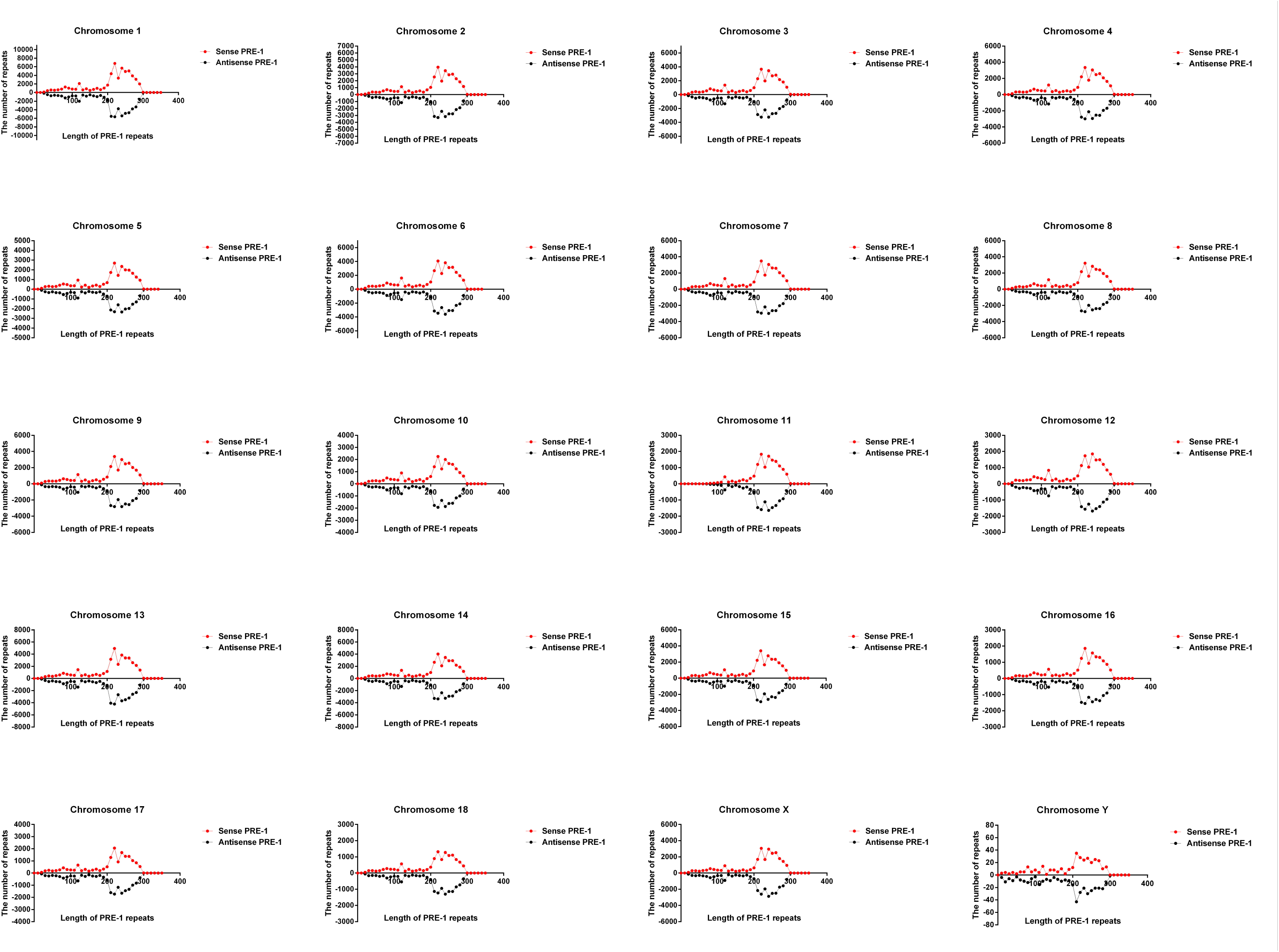
Length frequency distribution of PRE-1 elements on both strands of all 20 chromosomes on the pig.

**Fig. 3.**
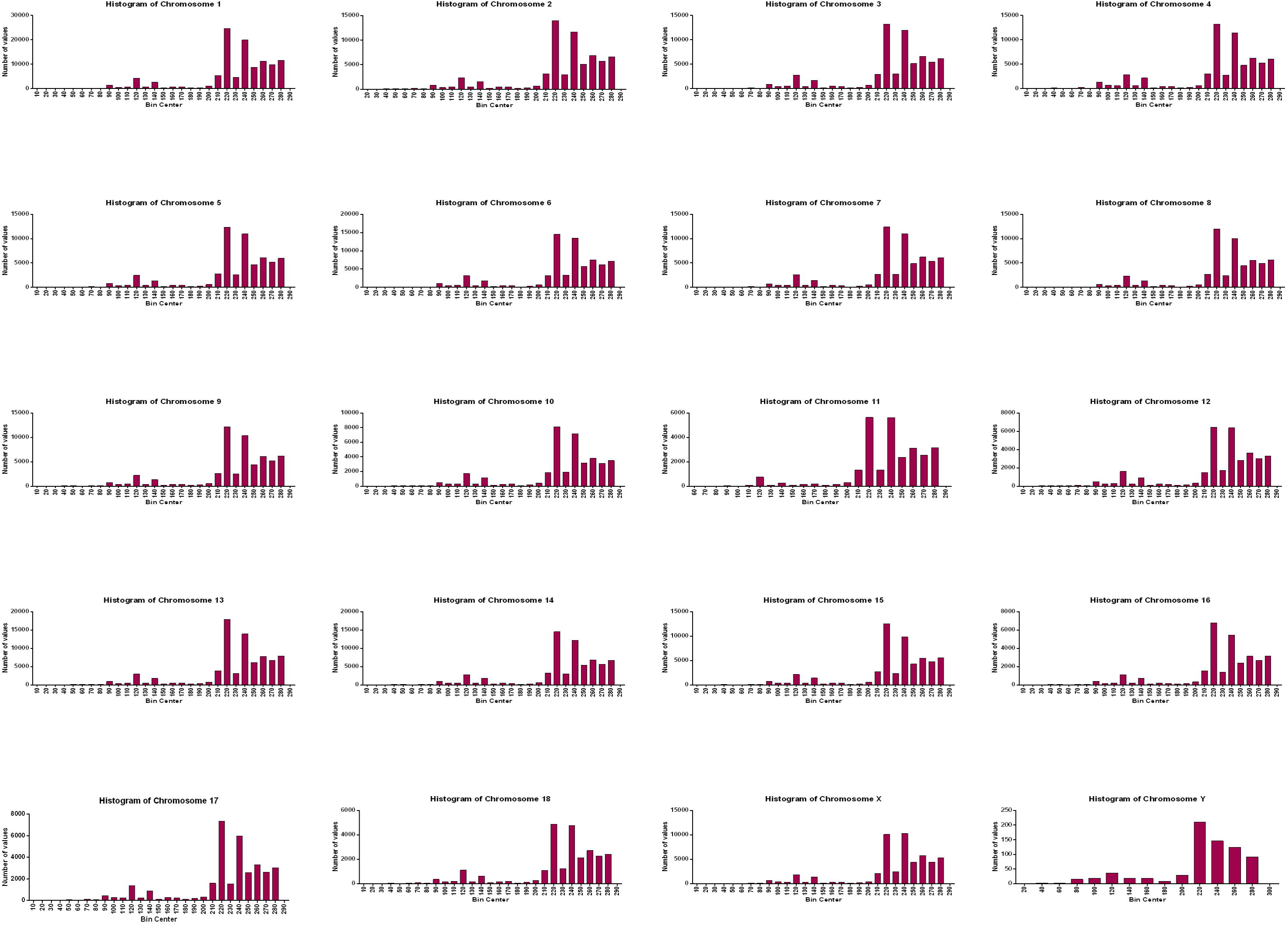
Fracture site frequency distribution of PRE-1 elements on all 20 chromosomes of the pig.

**Fig. 4.**
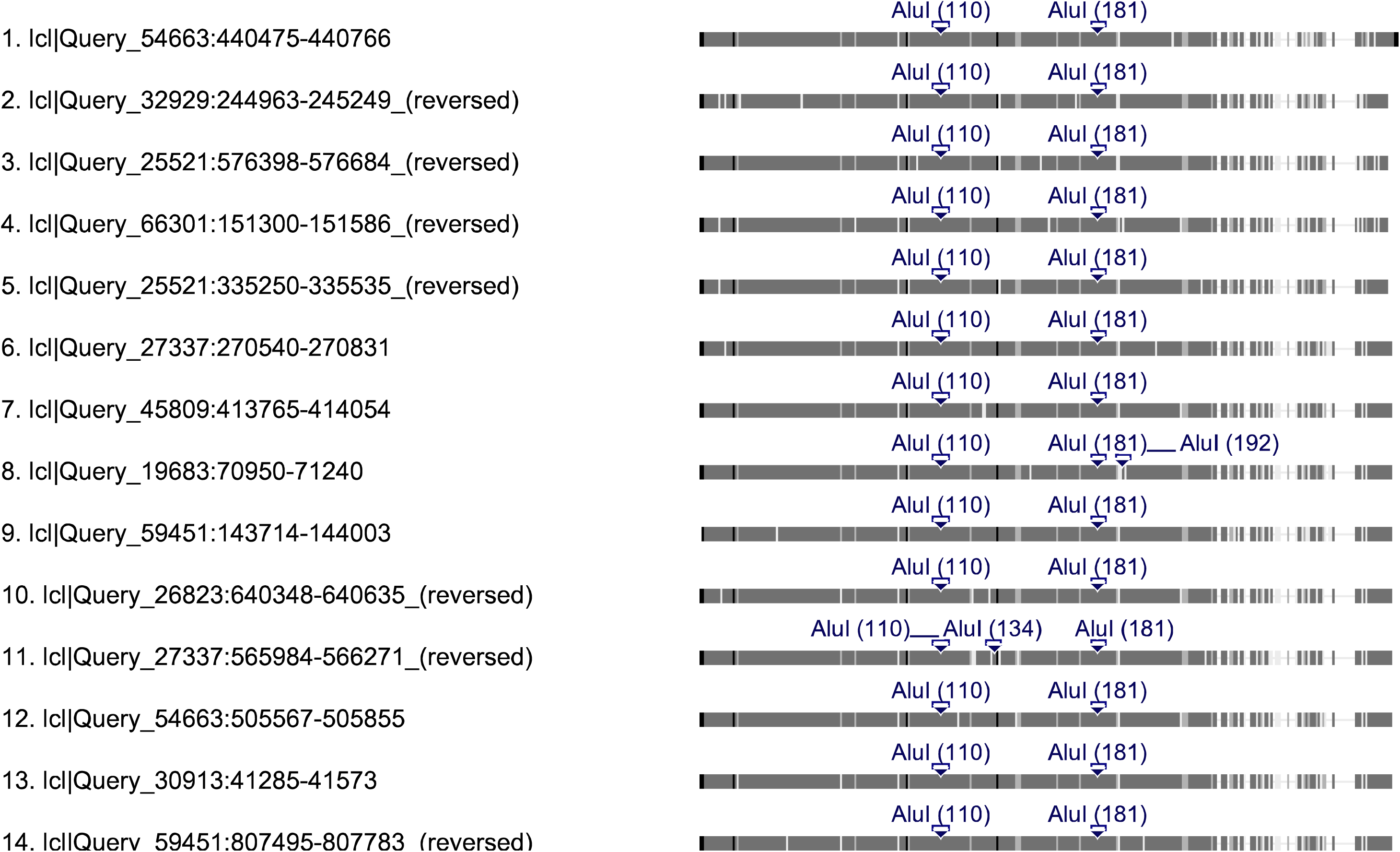
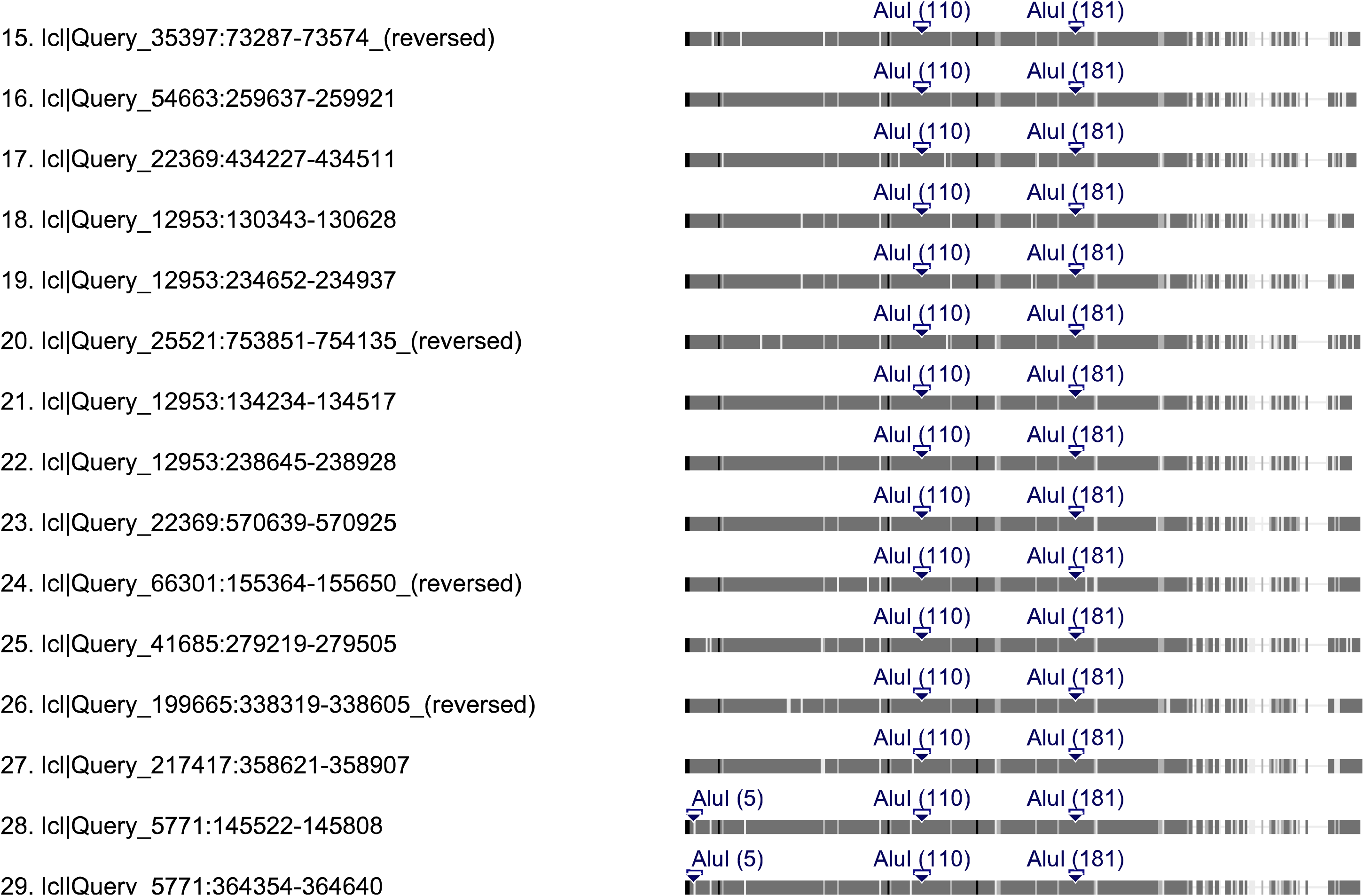
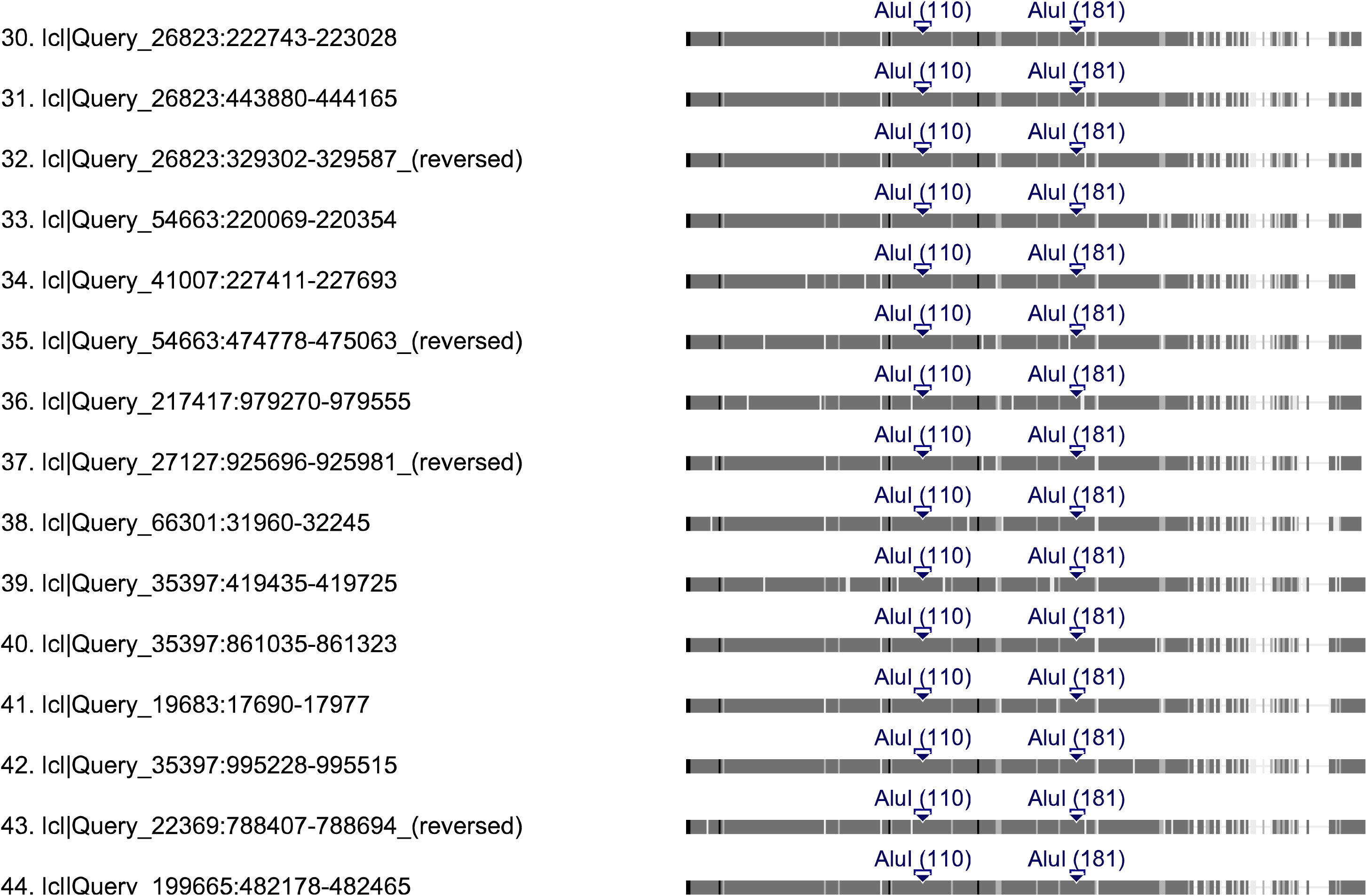
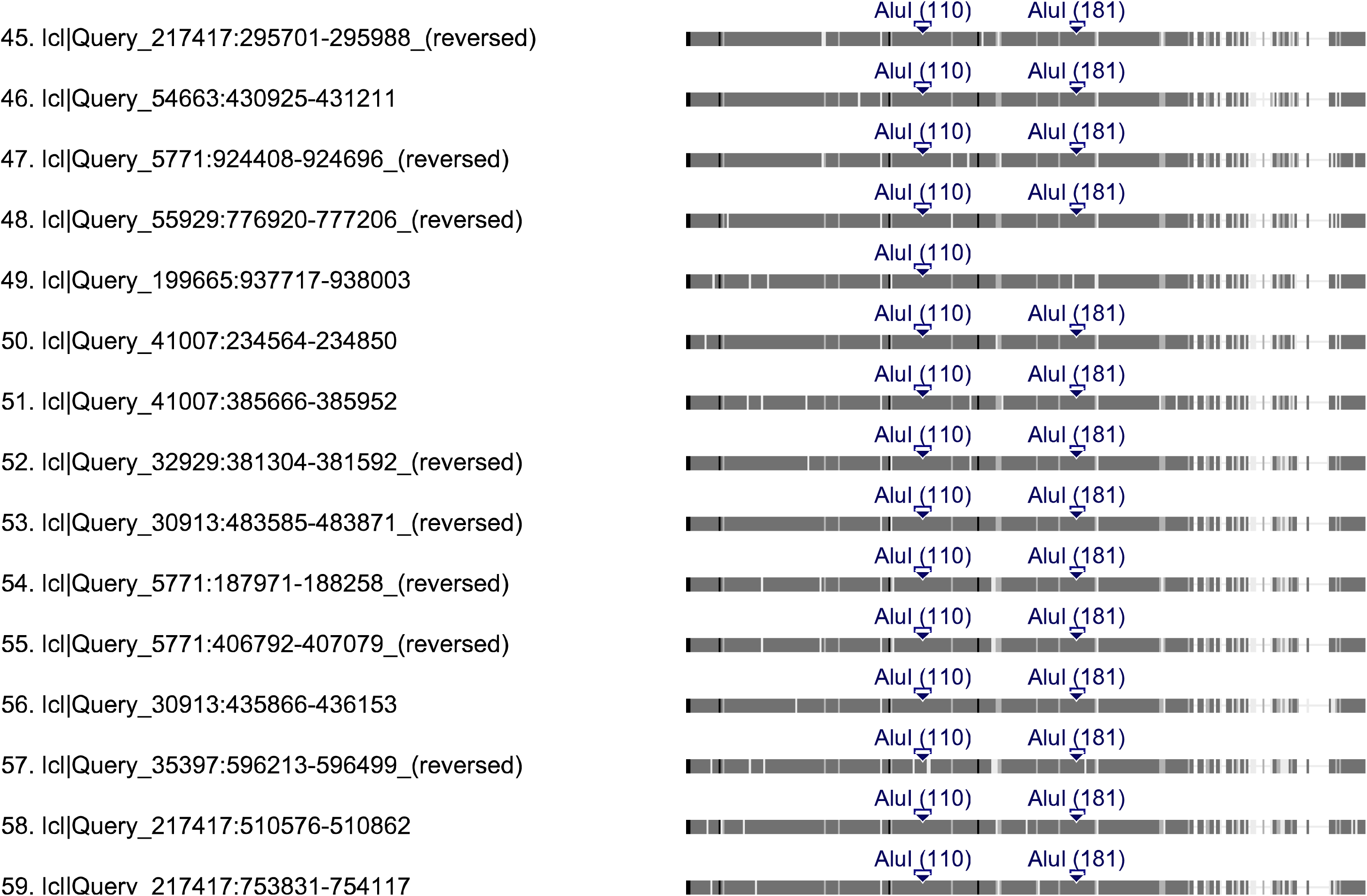
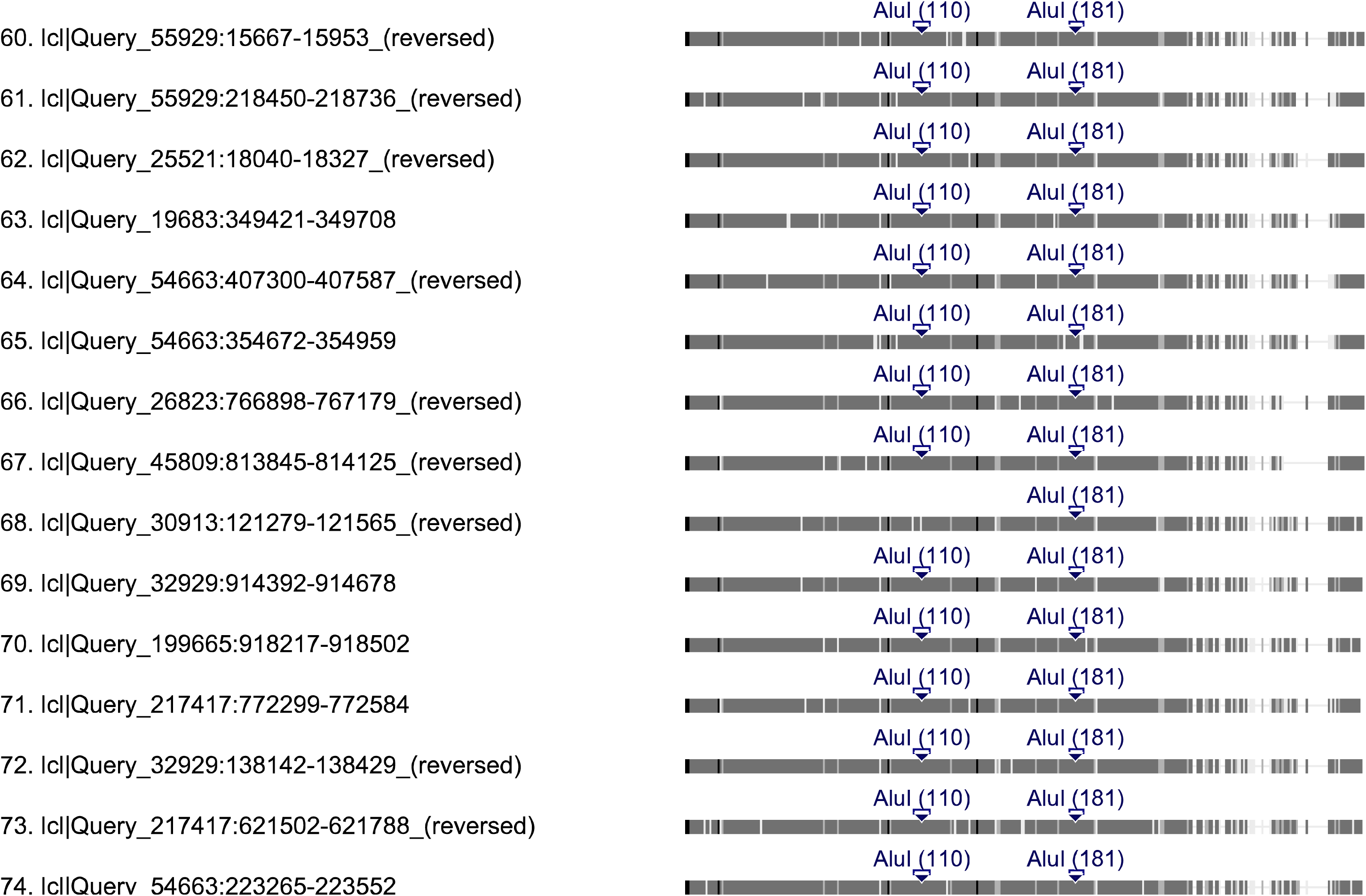
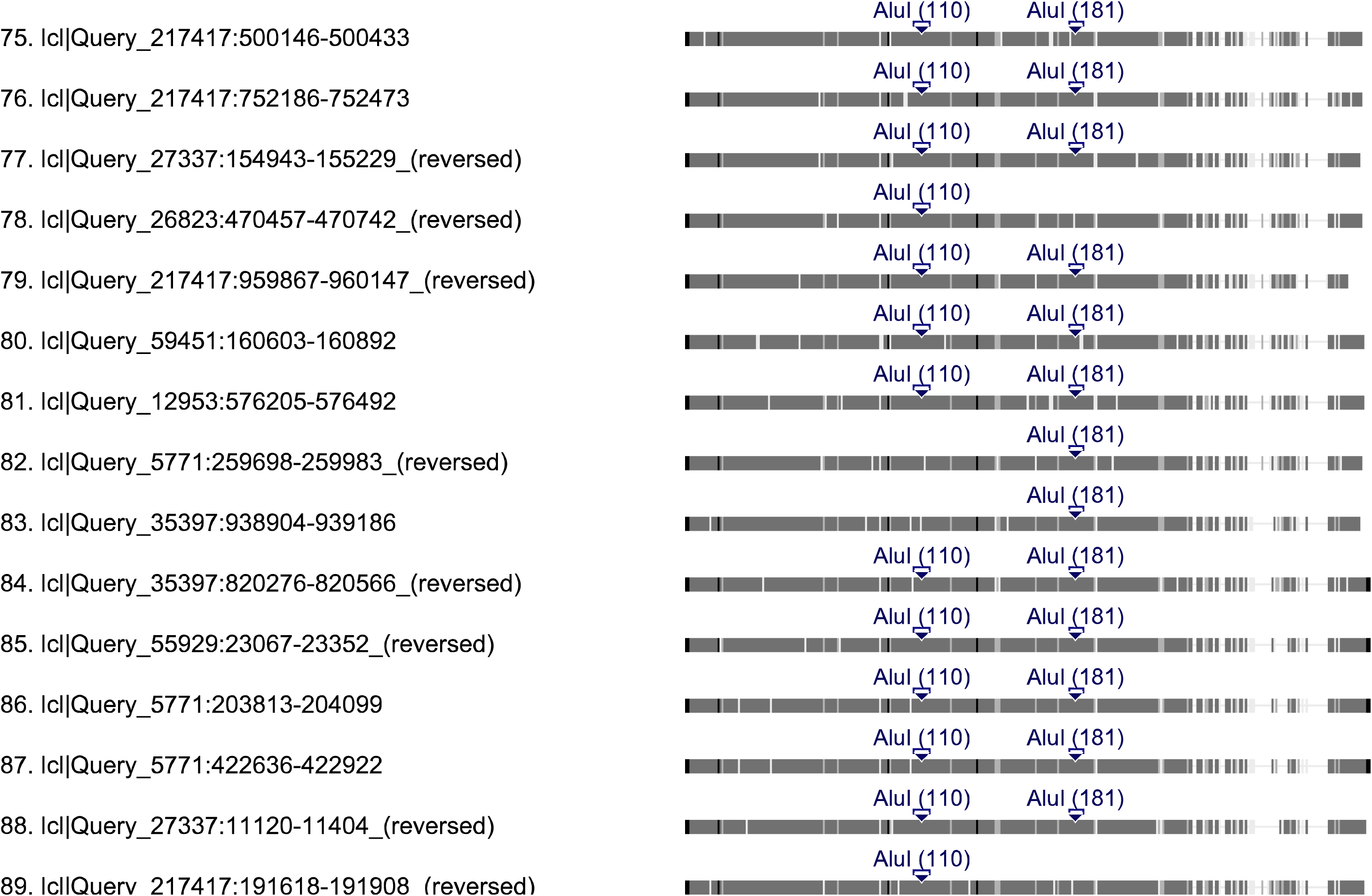
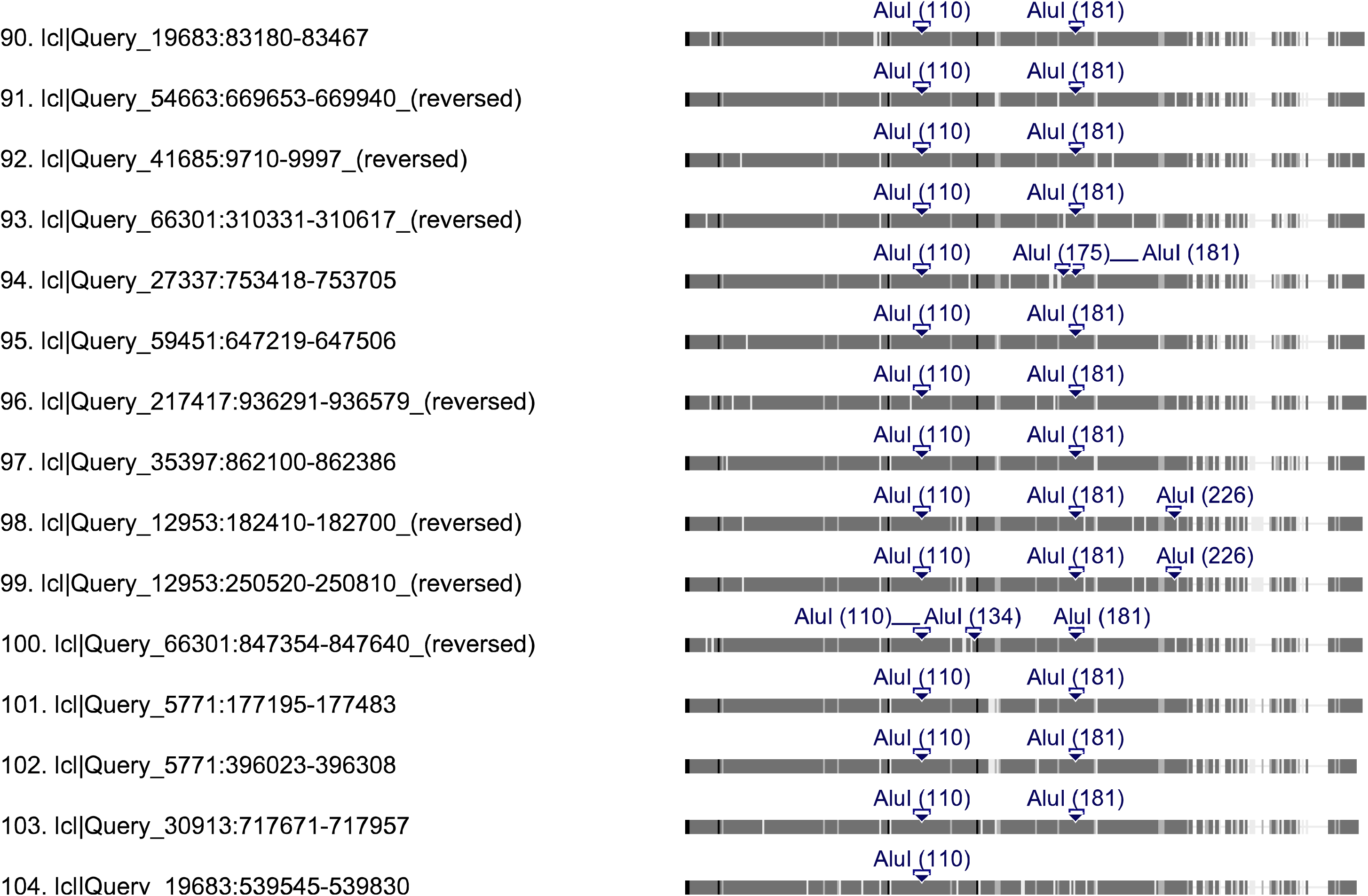
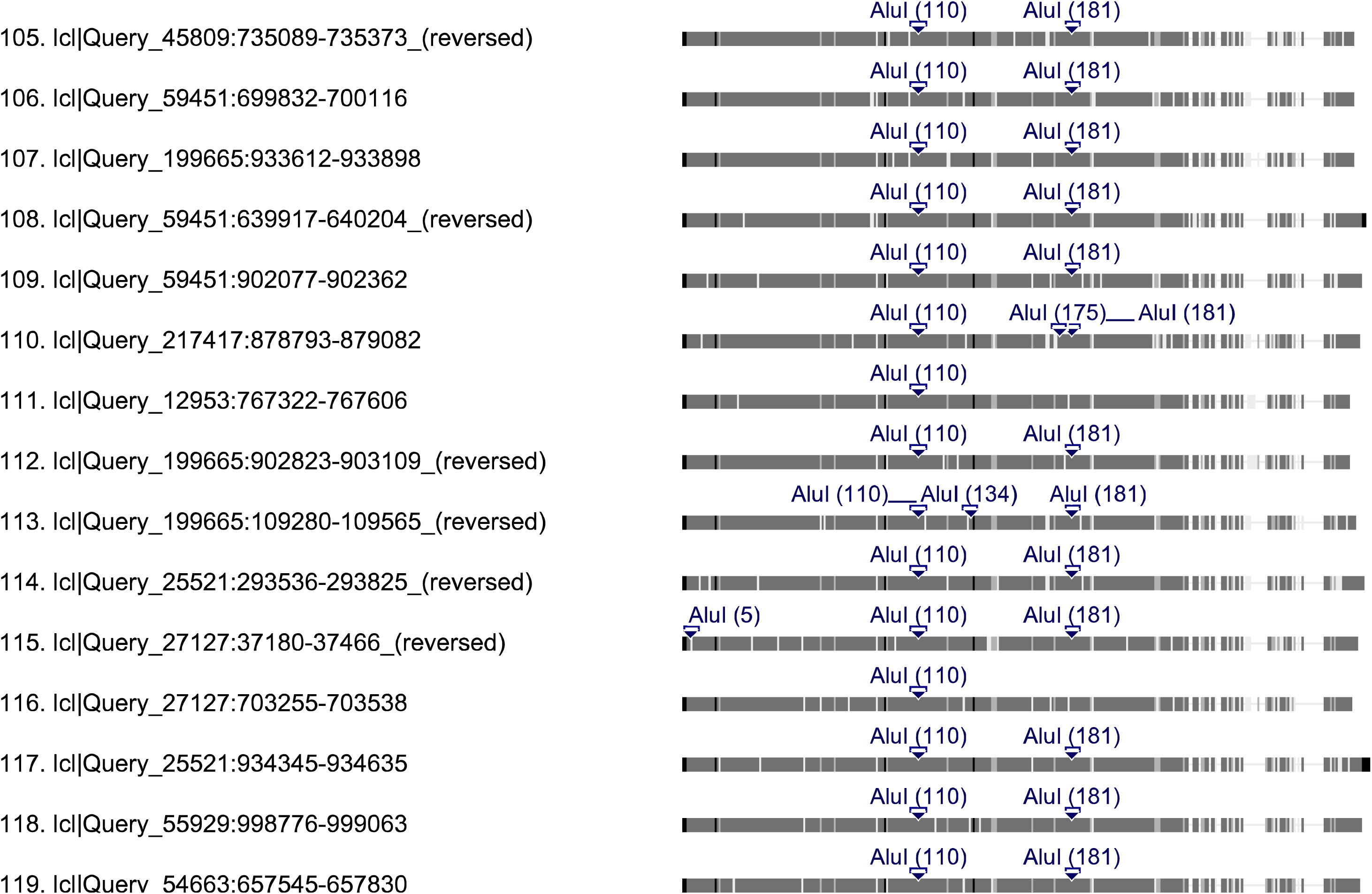
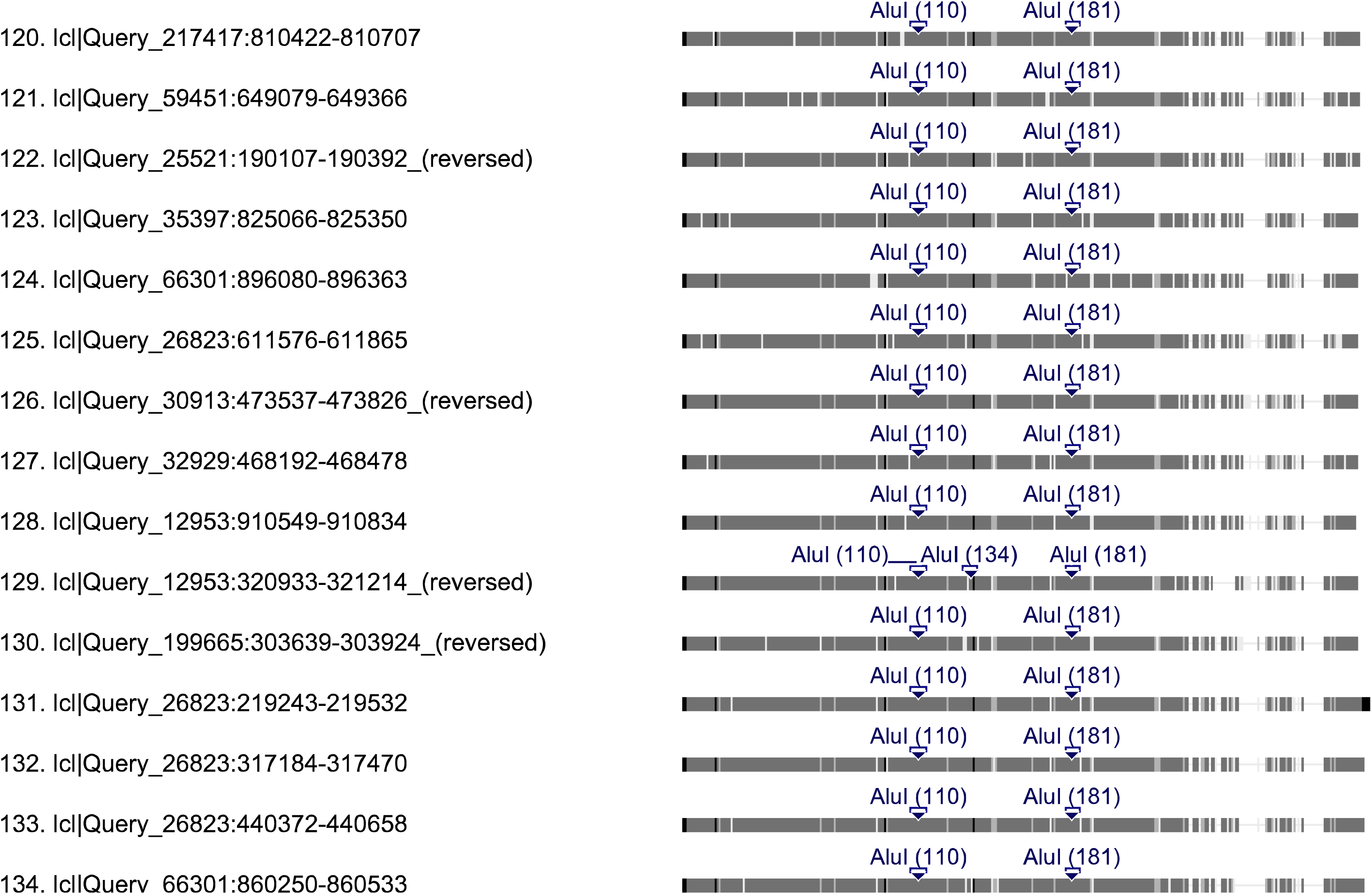
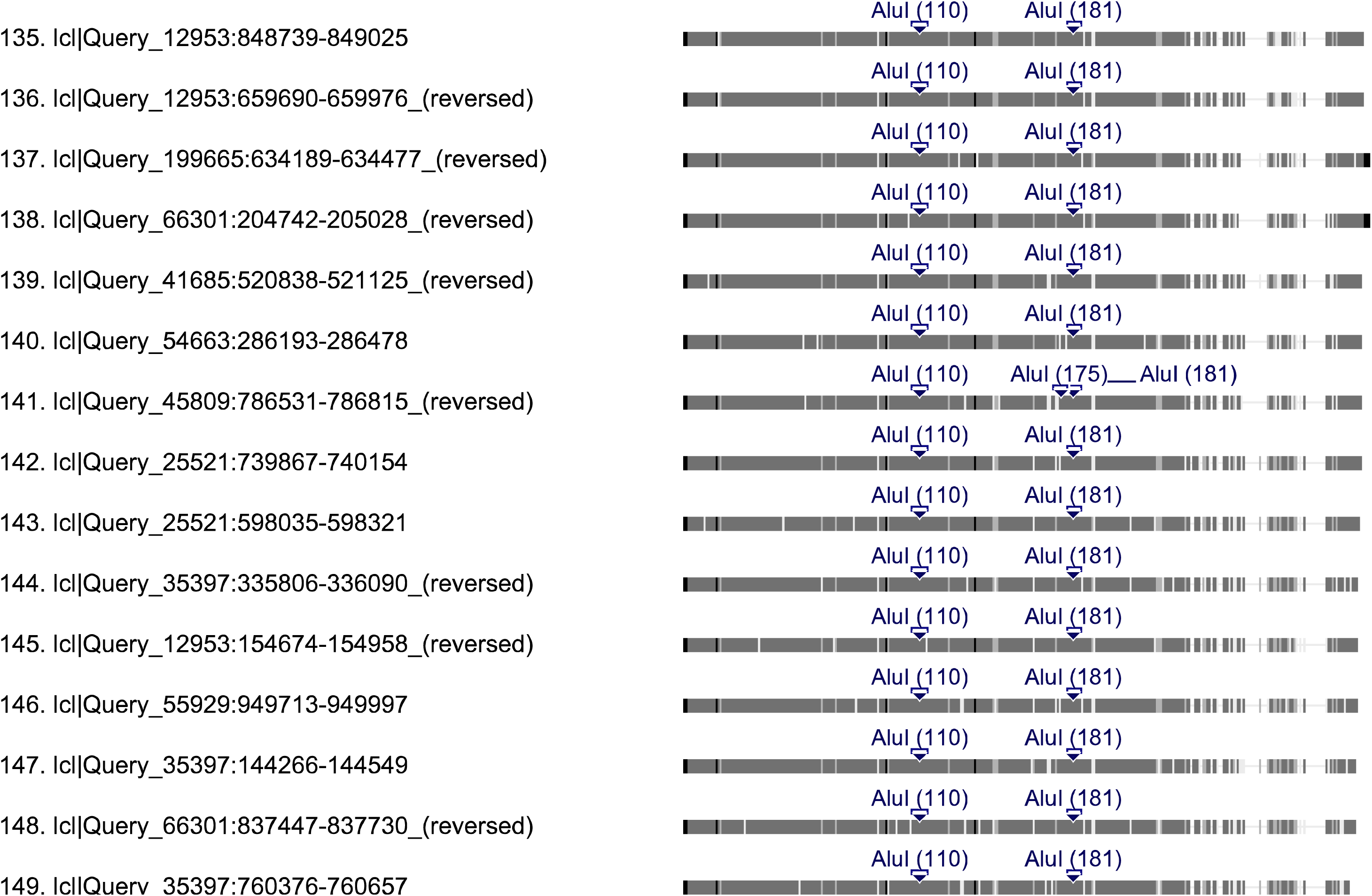
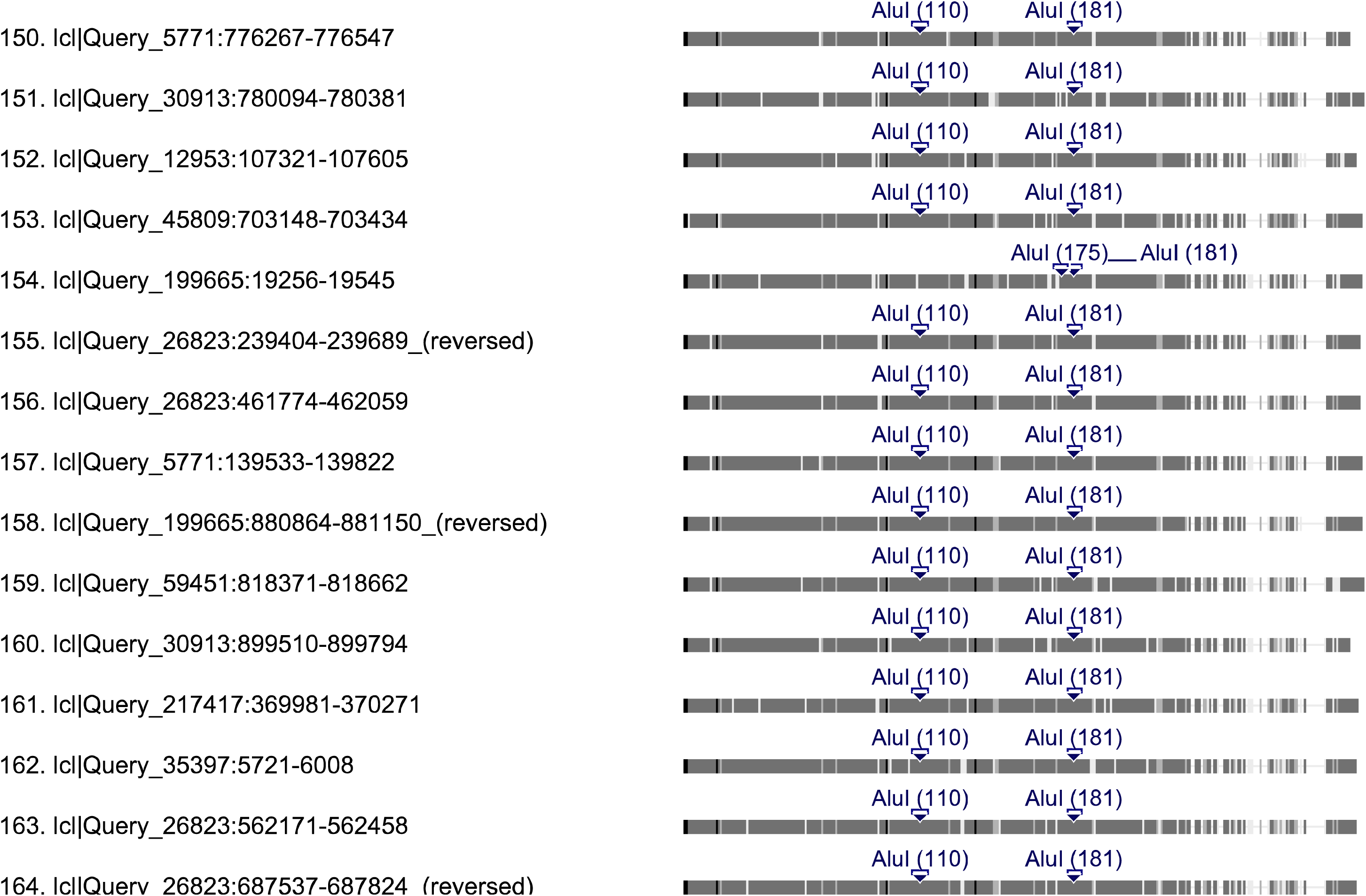
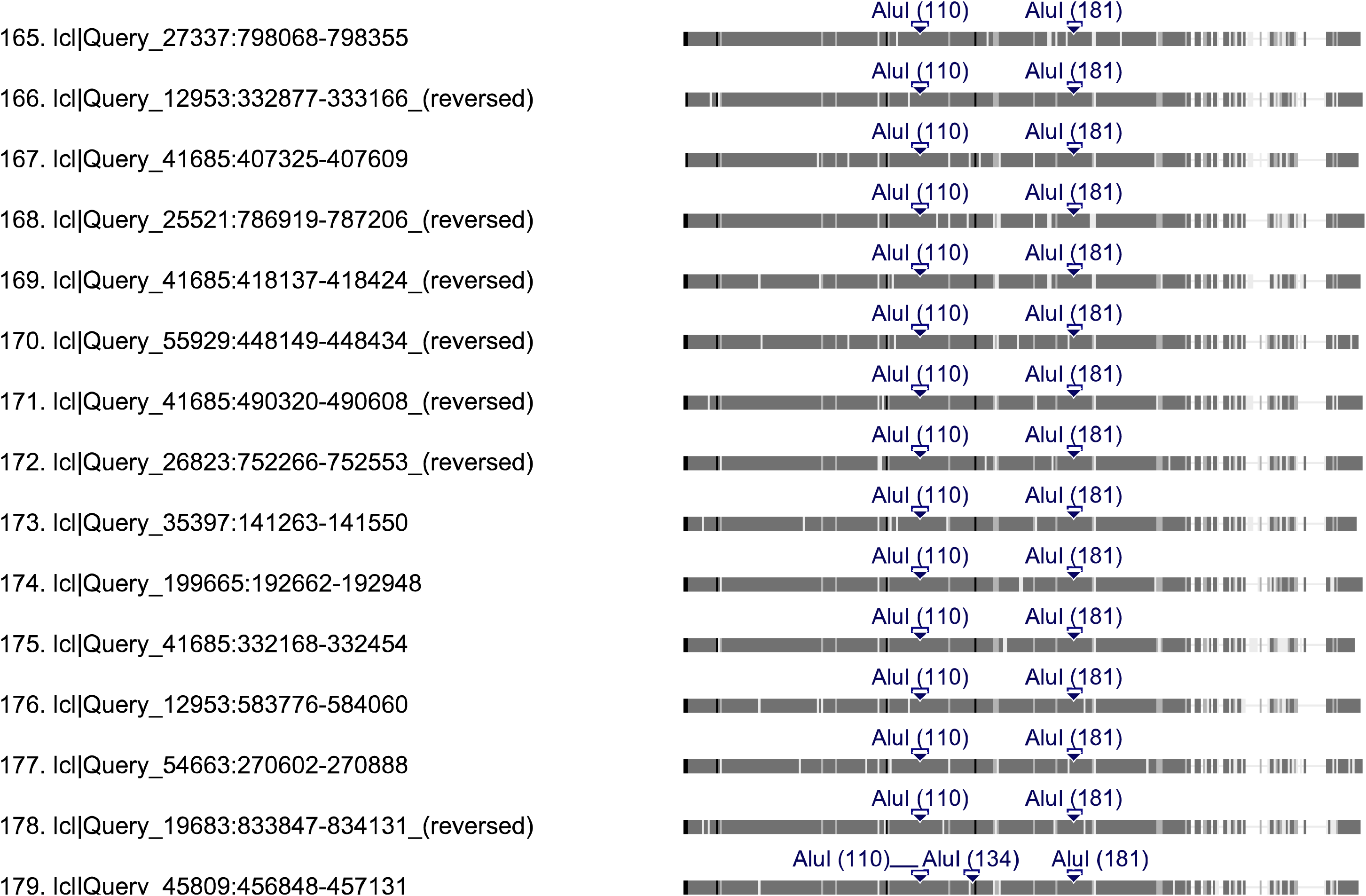
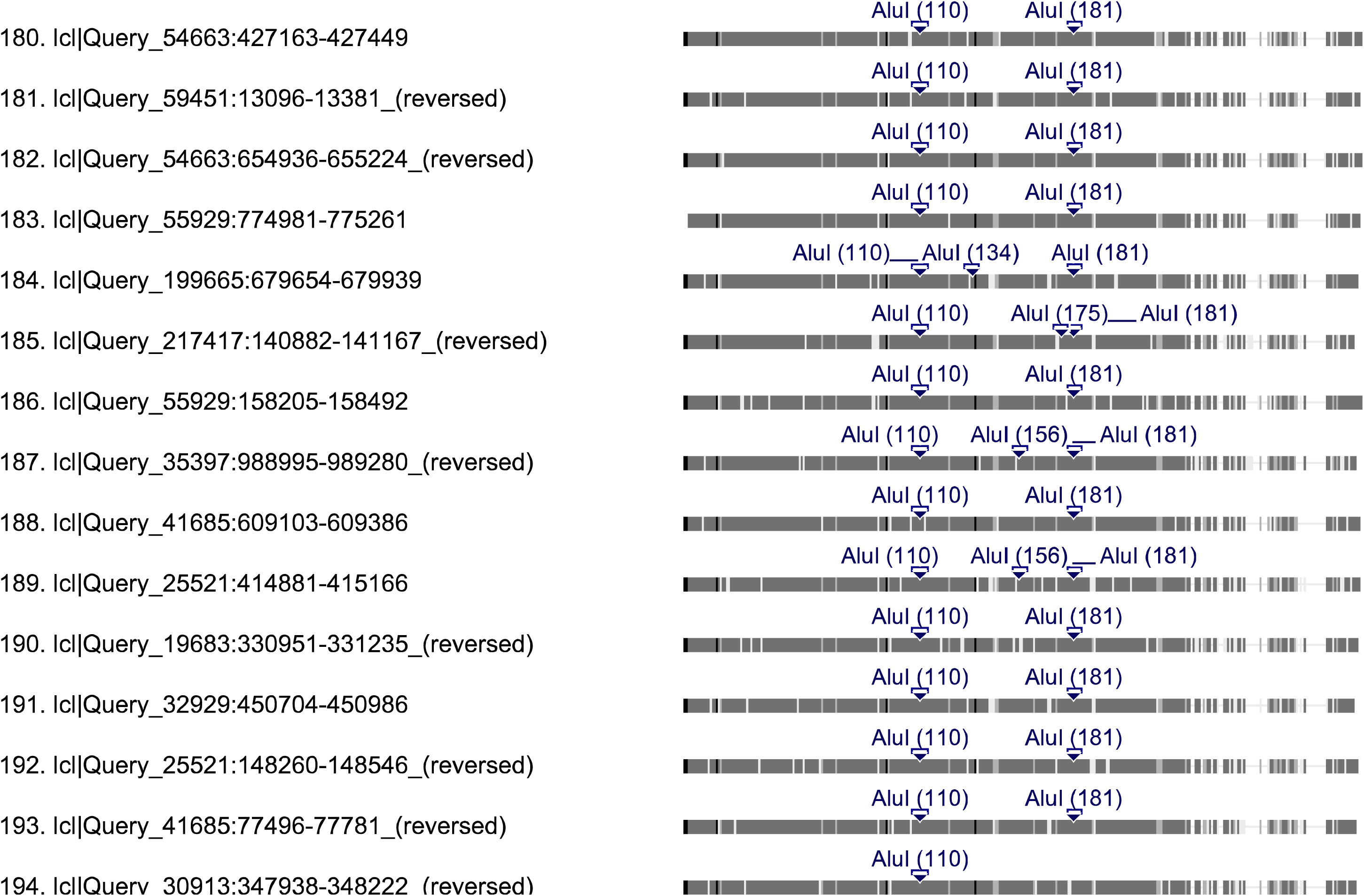
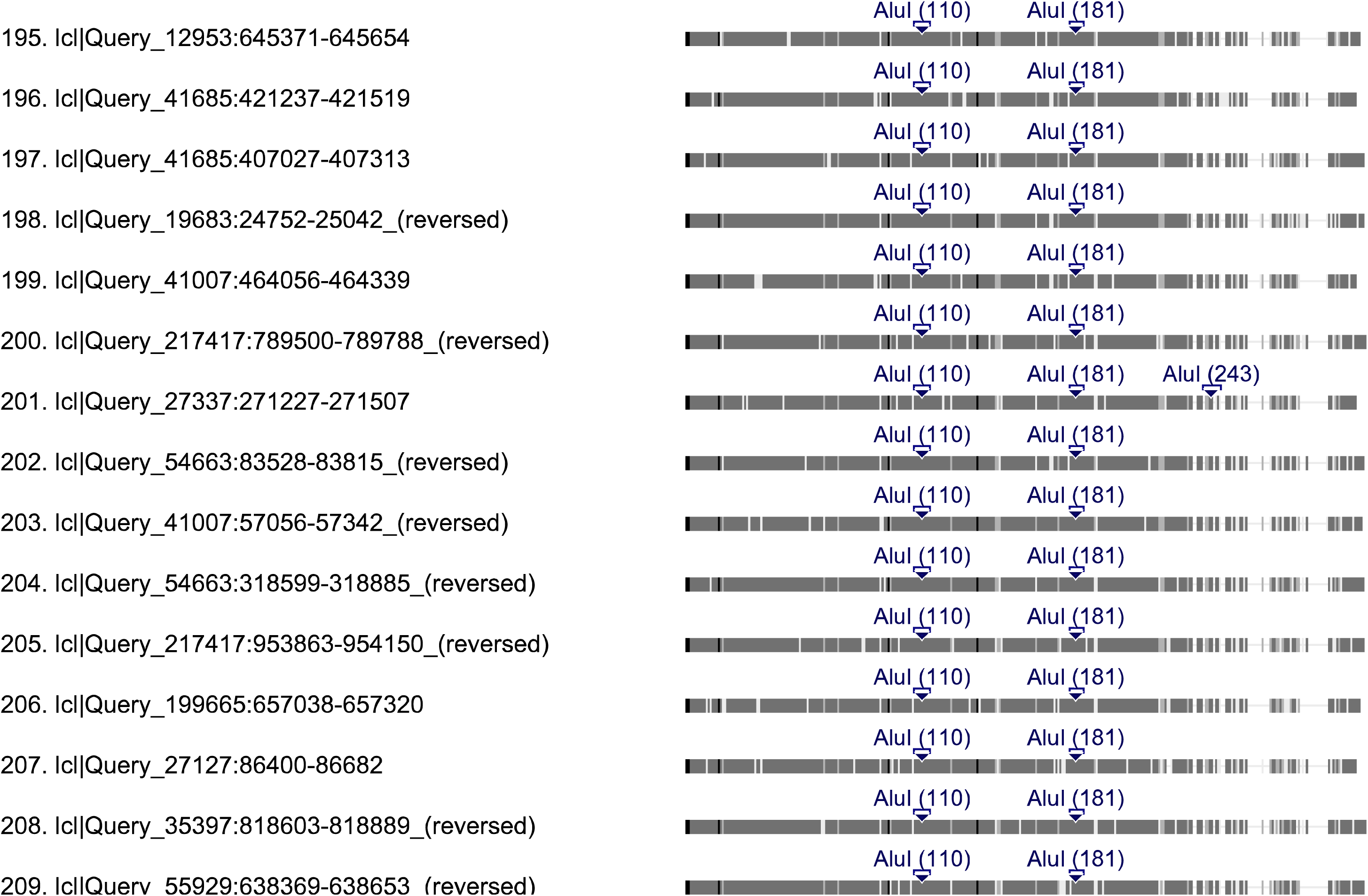
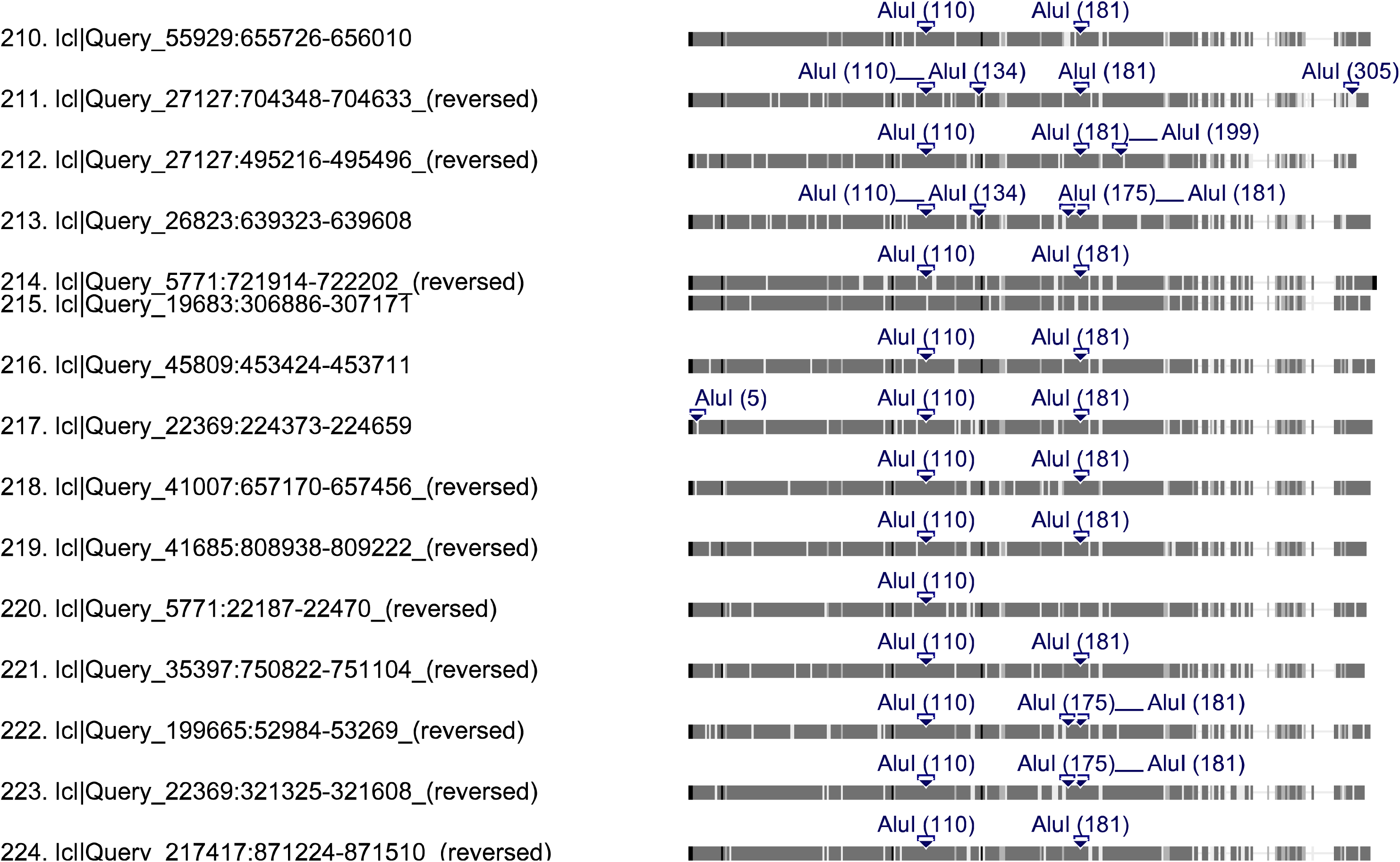
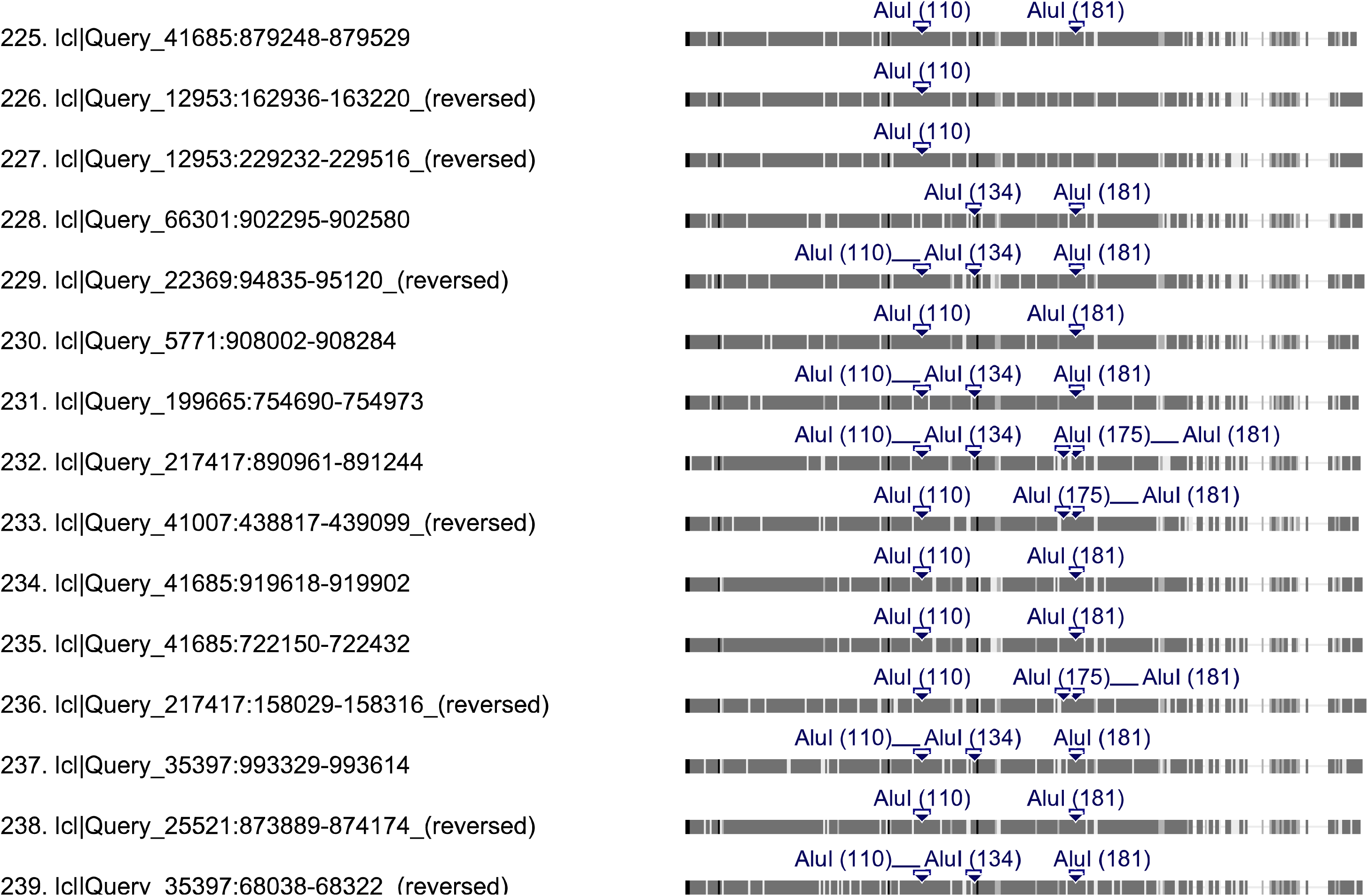
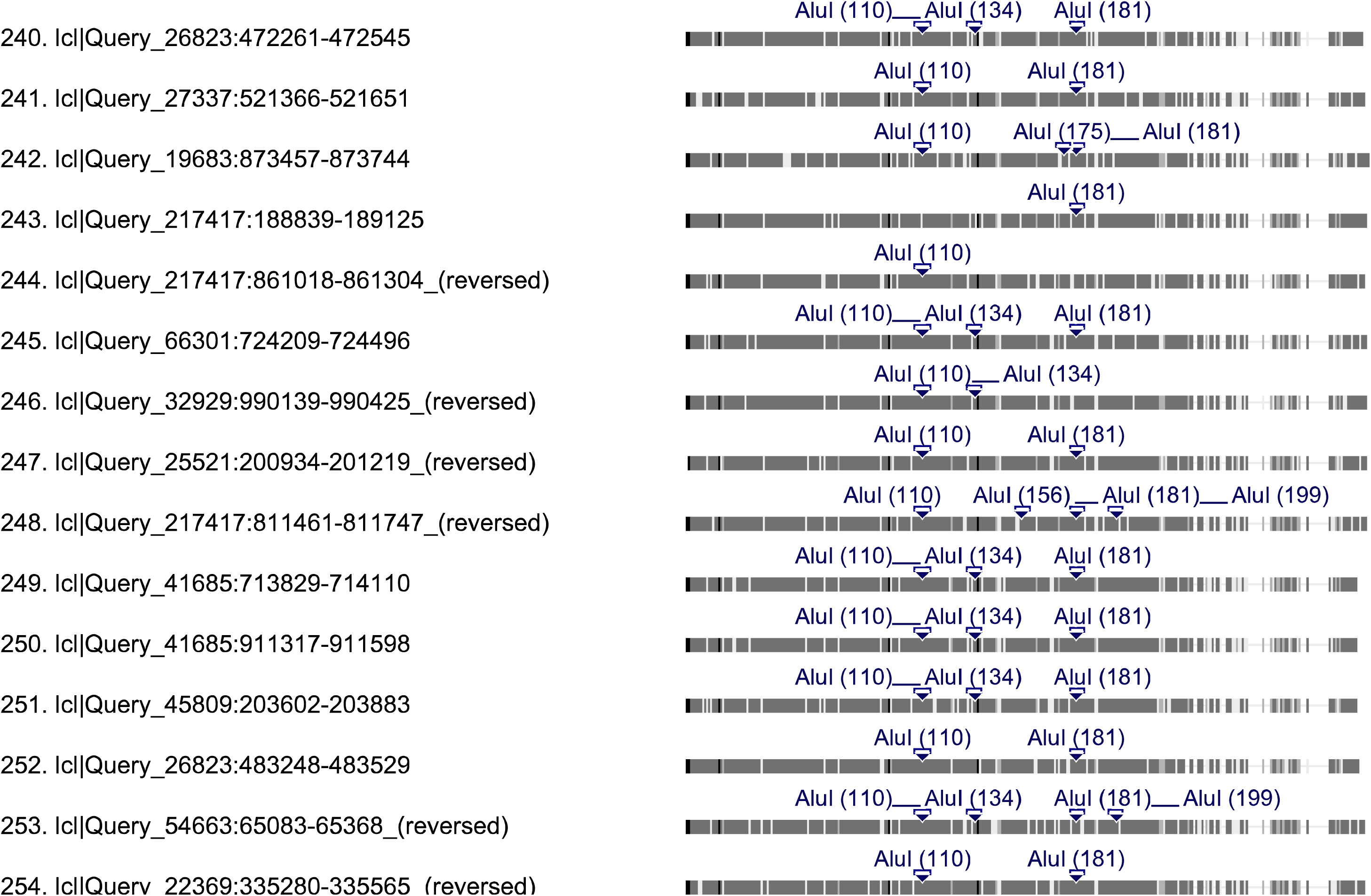
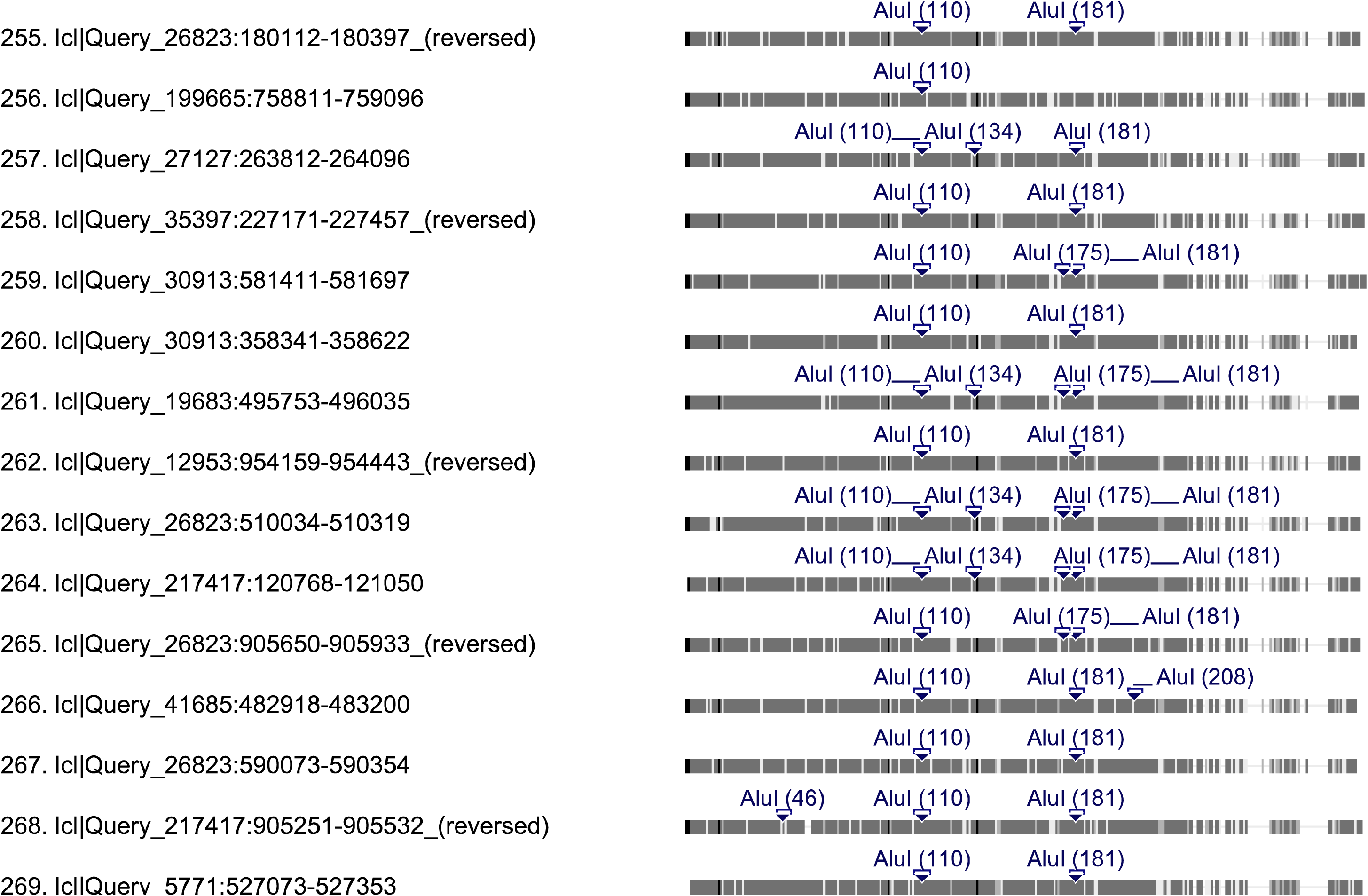
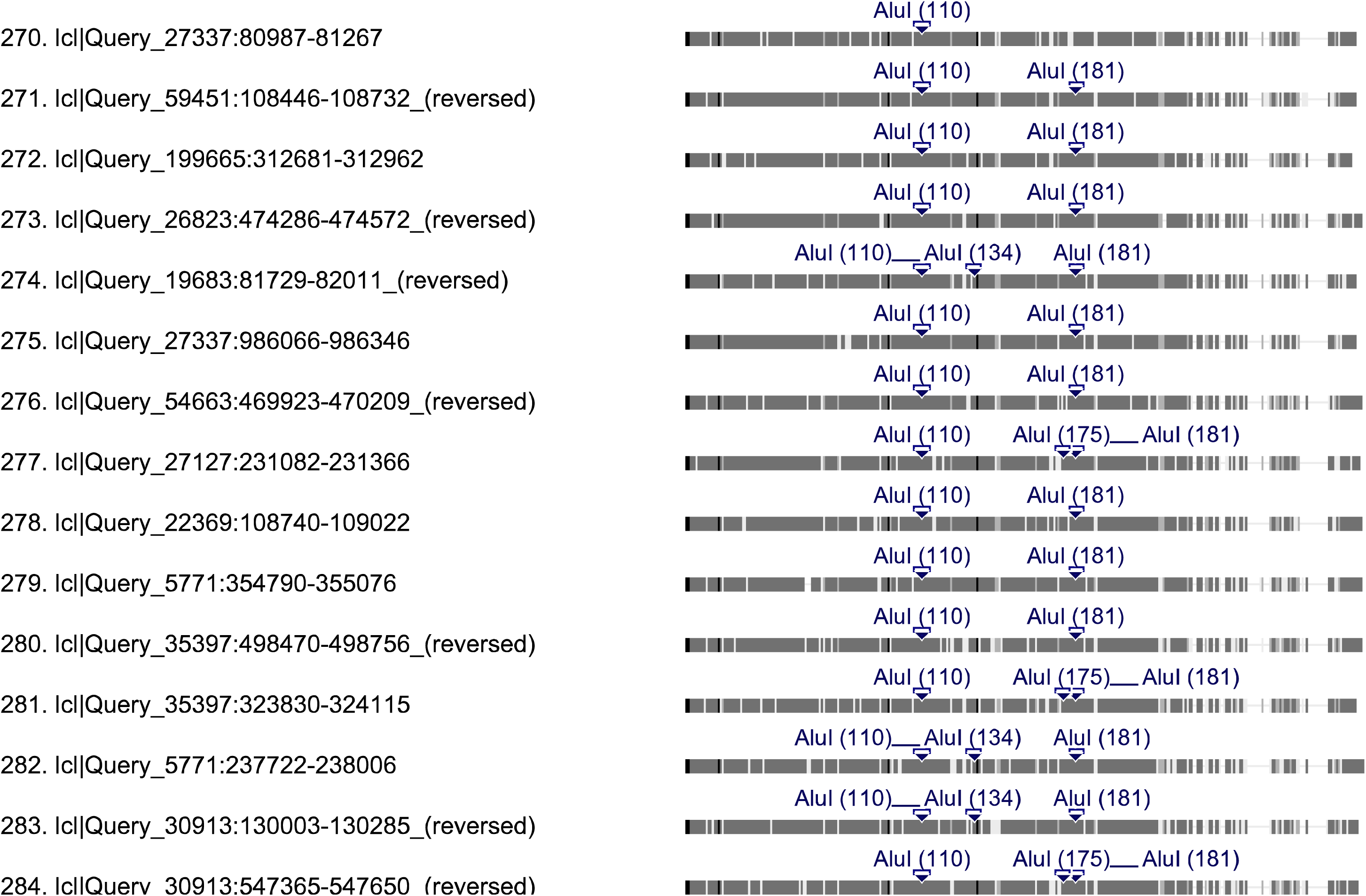
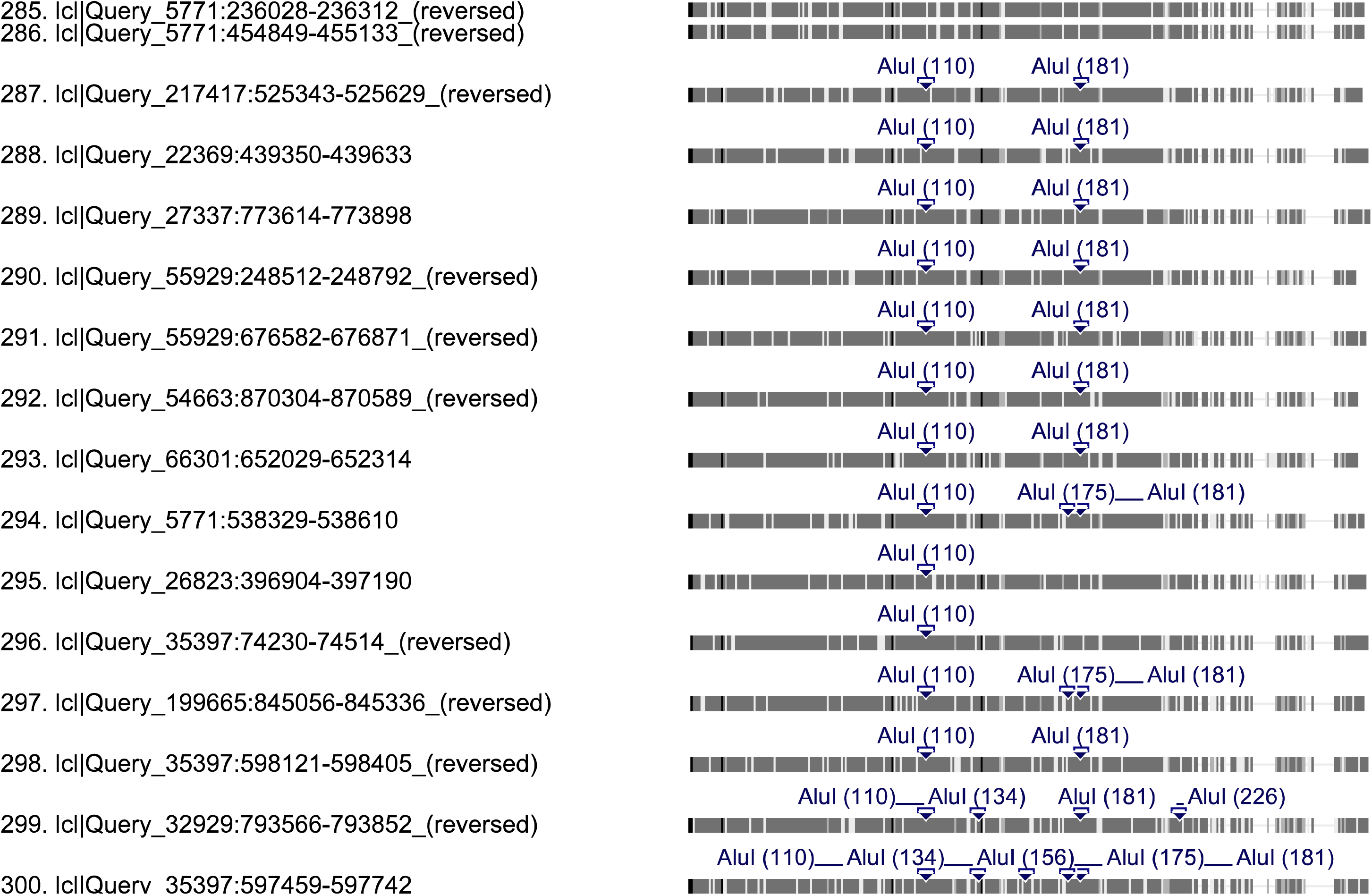
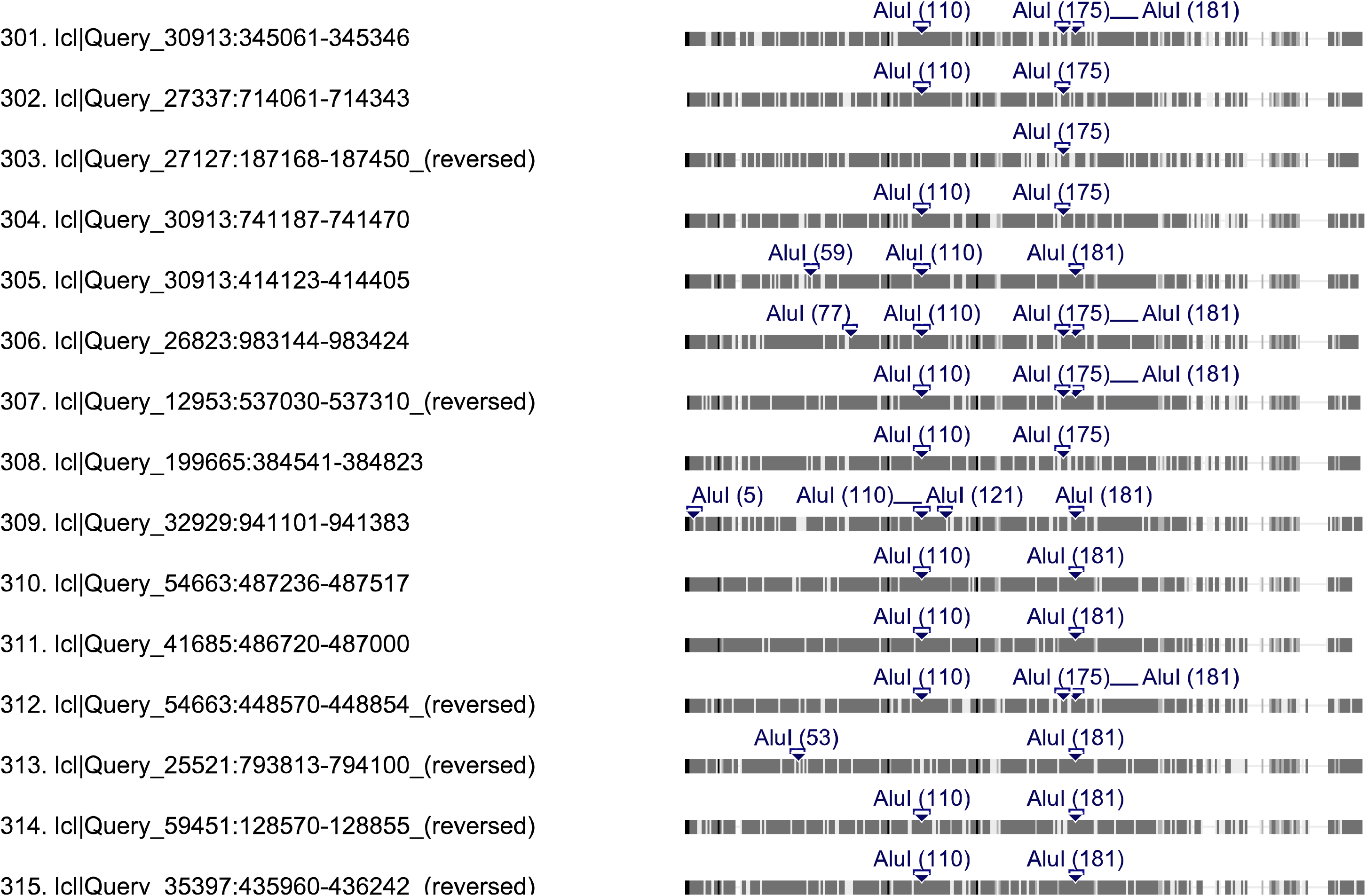
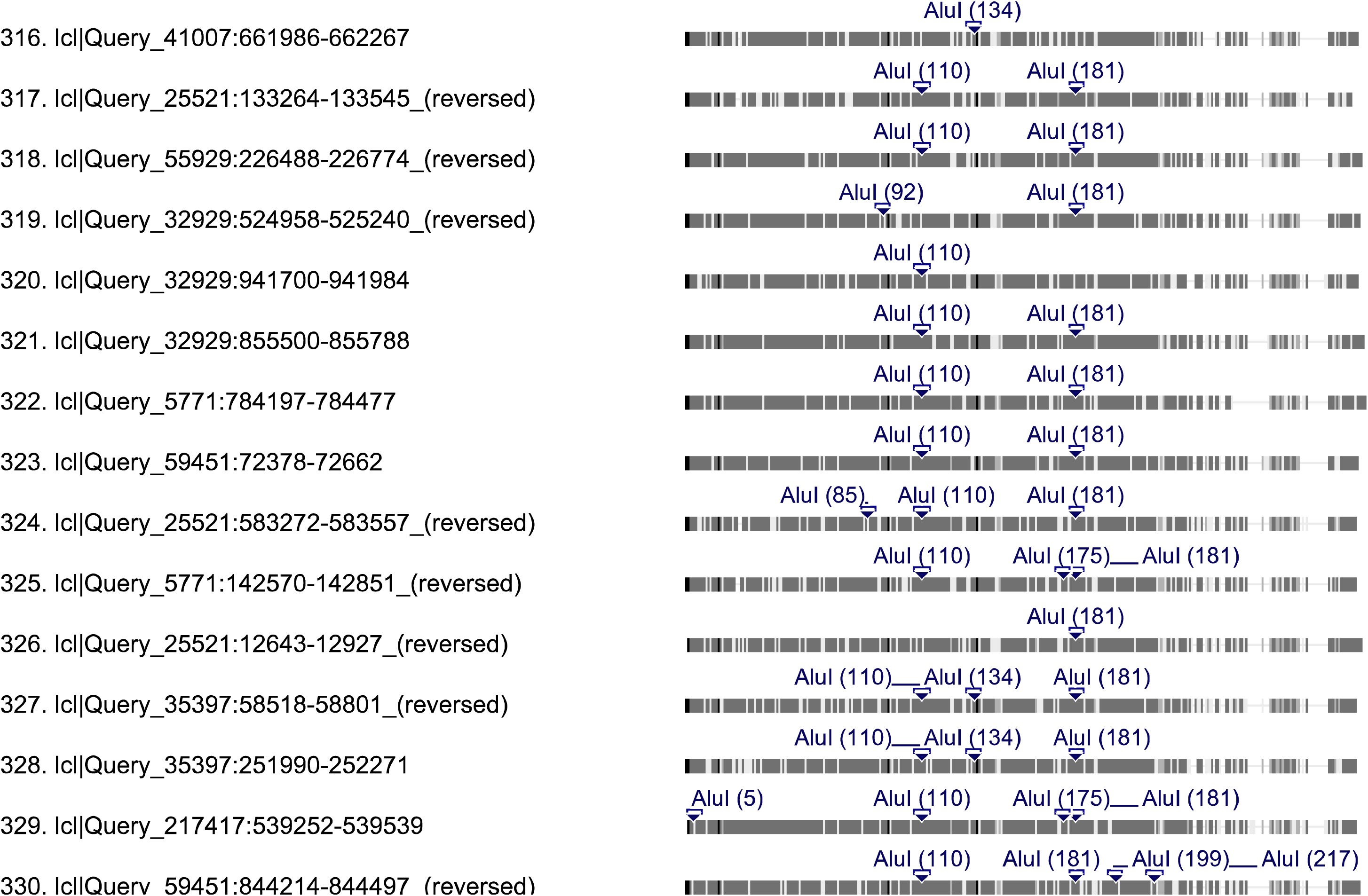
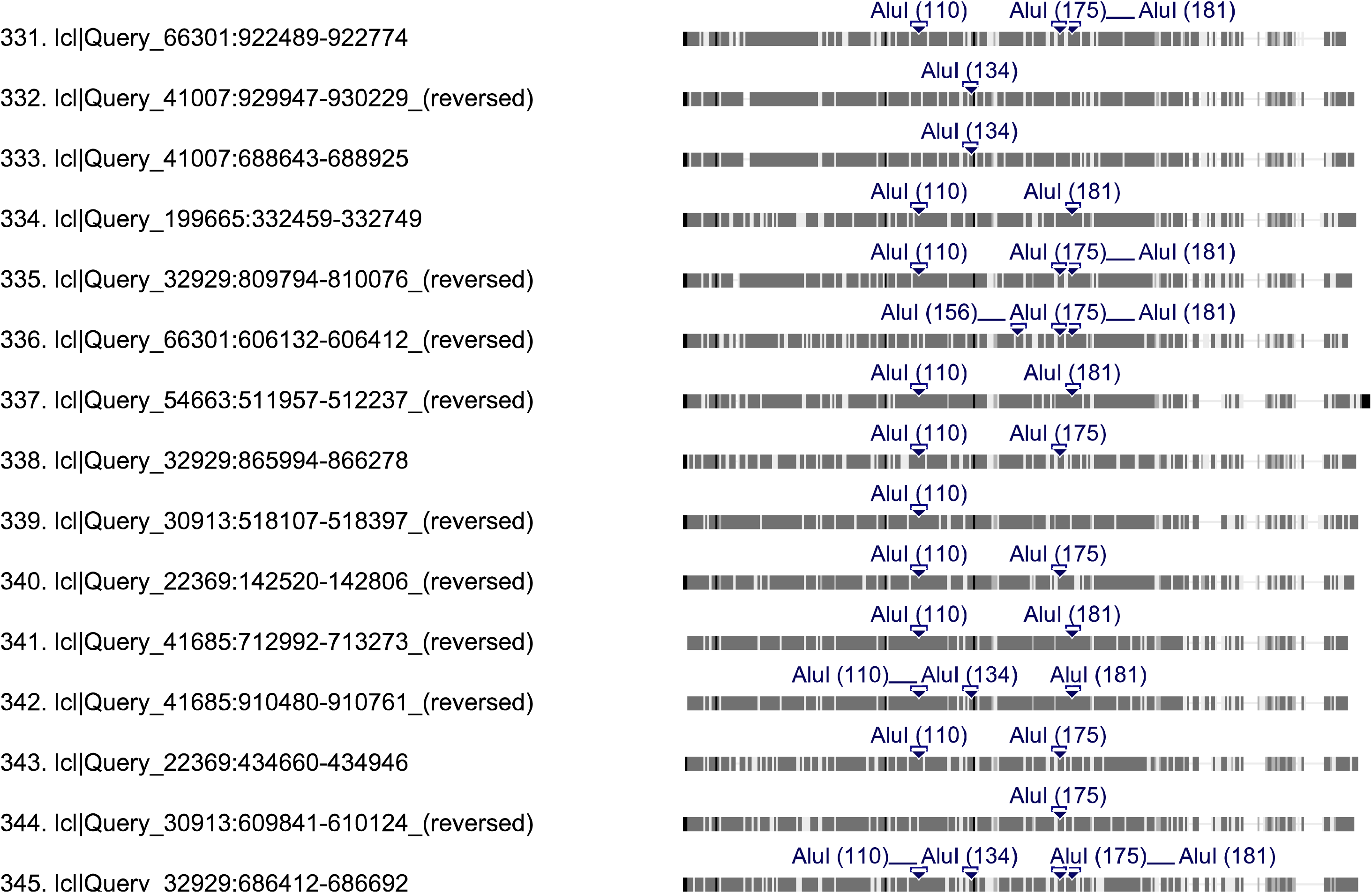
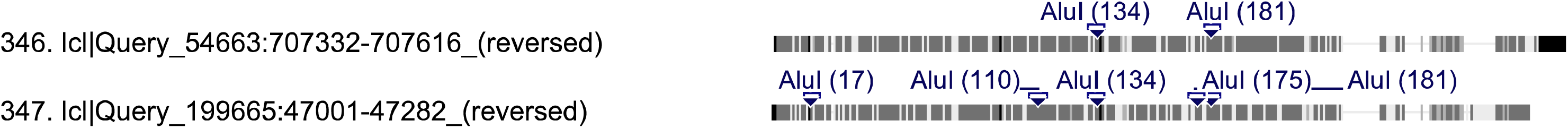
Distribution of AluI restriction enzyme recognition sites (AG^CT) on 347 PRE-1 elements.

Table 1 Information about genomic features for all 20 chromosomes of the pig.

A table describing the number of PRE-1 repeats, genes, protein-coding sequences, alternative splicing, exons, introns, and chromosome length of each pig chromosome.

Table 2 Information of the gene with the most exons on all 20 chromosomes of pig.

A table describing the number of PRE-1 repeats and exons in the gene with the most exons on each pig chromosome.

Table 3 Correlation coefficient distribution of density between PRE-1 repeats and genes on all 20 chromosomes of the pig.

A table describing the number of PRE-1 repeats and genes in interval lengths on each pig chromosome. Because the length and gene cluster of every chromosome isn’t same, every chromosome has its own optimal bin width value for its highest correlation coefficient. Through a series of calculations, we find that the bin width value of 10000000 or 5000000 is an ideal option to obtain a relatively high correlation coefficients, except for chromosome Y (bin width value is 200000), every correlation coefficient is chosen from the higher value between the frequency distribution of bin width values of 10000000 and 5000000.

Table 4 Length-frequency of PRE-1 repeats on all 20 chromosomes of the pig.

A table describing the frequency value of PRE-1 repeats on both strands of each pig chromosome. The frequency value of repeats on antisense strands are set negative.

Table 5 Paired *T*-test of correlation coefficients between sense/sense PRE-1 and antisense/antisense PRE-1 with sense/antisense PRE-1 on all 20 chromosomes of the pig.

A table, including correlation coefficient matrices, of sense PRE-1 and antisense PRE-1 among all 20 chromosomes and correlation coefficient table of sense and antisense PRE-1 of each chromosome of the pig. The correlation coefficient matrix was used to calculate the average correlation coefficient of each chromosome, and compared with that of sense and antisense PRE-1 of each chromosome to perform a paired T-test.

Table 6 Growth curve of the Pearson coefficient of the length frequency in the region ranging from 1 to 20000000 bp on chromosome 1.

A table describing the increase in the Pearson coefficient of the length frequency between repeats on both strands with the range increased, including the number of the various sizes of sense and antisense repeats in a gradually increasing range.

Table 7 Proportion of PRE-1 hits occupying the total transcribed RNAs

A table, including the raw data, for separating the exonized PRE-1 elements in the transcribed RNAs.

